# Global invasion history of the world’s most abundant pest butterfly: a citizen science population genomics study

**DOI:** 10.1101/506162

**Authors:** Sean F. Ryan, Eric Lombaert, Anne Espeset, Roger Vila, Gerard Talavera, Vlad Dincă, Mark A. Renshaw, Matthew W. Eng, Meredith M. Doellman, Emily A. Hornett, Yiyuan Li, Michael E. Pfrender, DeWayne Shoemaker

## Abstract

A major goal of invasion and climate change biology research is to understand the ecological and evolutionary responses of organisms to anthropogenic disturbance, especially over large spatial and temporal scales. One significant, and sometimes unattainable, challenge of these studies is garnering sufficient numbers of relevant specimens, especially for species spread across multiple continents. We developed a citizen science project, “Pieris Project”, to successfully amass thousands of specimens of the invasive agricultural pest *Pieris rapae*, the small cabbage white butterfly, from 32 countries worldwide. We then generated and analyzed genomic (ddRAD) and mitochondrial DNA sequence data for these samples to reconstruct and compare different global invasion history scenarios. Our results bolster historical accounts of the global spread and timing of *P. rapae* introductions. The spread of *P. rapae* over the last ∼160 years followed a linear series of at least four founding events, with each introduced population serving as the source for the next. We provide the first molecular evidence supporting the hypothesis that the ongoing divergence of the European and Asian subspecies of *P. rapae* (∼1,200 yrBP) coincides with the domestication of brassicaceous crops. Finally, the international success of the Pieris Project allowed us to nearly double the geographic scope of our sampling (i.e., add >1,000 specimens from 13 countries), demonstrating the power of the public to aid scientists in collections-based research addressing important questions in ecology and evolutionary biology.

**Non-technical summary:** We provide genetic evidence that the success of the small cabbage white butterfly—its rise to one of the most widespread and abundant butterflies on the planet— was largely facilitated by human activities, through the domestication of its food plants and the accidental movement of the butterfly by means of trade and human movement (migration). Through an international citizen science project—Pieris Project—people from around the world helped to unravel the global invasion history of this agricultural pest butterfly by collecting samples for DNA analysis. The success of this citizen science project demonstrates the power of the public to aid in collections-based research that address important questions related to ecology and evolutionary biology.

## Introduction

Invasive species—species spread to places beyond their natural range, where they generate a negative impact (e.g., extirpate or displace native fauna, spread disease, destroy agricultural crops^1^)—continue to increase in number, with no signs of saturation^2^. The spread of invasive species often is driven by (human) migration, global trade and transportation networks^3^, and, in some cases, domestication of wild plants and animals^4^. A critical, often first step to mitigating the spread and impacts of invasive species is to understand their invasion history, including assessing source populations, routes of spread, number of independent invasions, and the effects of genetic bottlenecks, among other factors. Such detailed knowledge is crucial from an applied perspective (e.g., developing an effective biological control program) as well as for addressing basic questions associated with the invasion process (e.g., genetic changes and adaptation to novel environments)^5^.

Unraveling a species’ invasion history often requires sampling across large spatial and temporal scales, which can be challenging and expensive, particularly for many invasive species found on multiple continents. Citizen science—research in which non-scientists play a role in project development, data collection or discovery and is subject to the same system of peer review as conventional science^6^—is a potentially powerful means to overcome some of these challenges. A major strength of citizen science is that it can greatly enhance the scale and scope of science and its impact on society^7^. Consequently, there are now thousands of citizen science projects worldwide. Yet, still very few involve agricultural pests^6^ and nearly all rely on observations (e.g., sightings or photographs), limiting their capacity to address fundamental questions in ecology and evolution.

*Pieris rapae*, the small cabbage white butterfly, is the world’s most widespread and abundant pest butterfly. Caterpillars of this species are a serious agricultural pest of crops in the Brassicaceae family (e.g., cabbage, kale, broccoli, brussels sprouts)^8^. This butterfly is believed to have originated in Europe and subsequently undergone a range expansion into Asia several thousand years ago as a result of domestication and trade of its host plants^9,10^. The Europe and Asia populations recognized today are believed to represent separate subspecies—*P. rapae rapae* and *P. rapae crucivora*, respectively.

The small cabbage white butterfly has been introduced to many other parts of the world over the last ∼160 years. These invasions are unique in that there is a wealth of historical records (observations and collections) documenting the putative dates of first introduction (North America in the 1860s^11^, New Zealand in 1930^12^, and Australia in 1937^13^). Detailed accounts and observations from what was essentially a 19^th^ century citizen science project led by the entomologist Samuel Scudder provide a chronology of the spread of *P. rapae* across North America and suggest that there were multiple independent introductions^11^. While the small cabbage white butterfly ranks as one of the most successful and abundant invasive species, a detailed analysis of its invasion history has never been undertaken^9,14^. In addition, the consequences of this rapid invasion on the population genetic structure and diversity also are unknown.

Here, we employ a collection-based citizen science approach to obtain range-wide, long-term, population-level sampling of this globally distributed invasive agricultural pest. Molecular genomics tools are then applied to this global collection of specimens to demonstrate that a citizen science approach can be used to address a multitude of important questions in ecology and evolutionary biology, including the reconstruction of the global invasion history of *P. rapae* and assessment of historical and contemporary patterns of genetic structure and diversity.

## Results

### Citizen scientist assisted sampling

The international citizen science project—Pieris Project—recruited more than 100 participants that collected >1,000 butterflies from 13 countries in less than three years. The majority of our participants (citizen scientists) were recruited through entomological and lepidopterist societies and other organizations related to nature and science. These citizen scientist collections were supplemented with collections from researchers bringing the total to >3,000 *P. rapae* from the period of 2002-2017 (median collection year: 2014, Fig S1). Nearly half (338/794) of the specimens used to generate mtDNA or ddRADseq data were from citizen scientists. Of the 32 countries represented in our collection, five countries (Portugal, Czech Republic, Gibraltar, Turkey, and South Korea) were made up of specimens sampled entirely by citizen scientists, three countries (Russia, Australia, and New Zealand) had the majority of specimens coming from citizen scientists, and three countries (United States, Canada, and Spain) had nearly half the specimens coming from citizen scientists (Table S1). These samples collectively cover nearly the entire native and invaded ranges, consisting of 293 localities spanning 32 countries (Fig 1, up-to-date collections map); note, we do not have collections from South America because there are no (known) populations of *Pieris rapae* on the continent.

**Fig 1.**
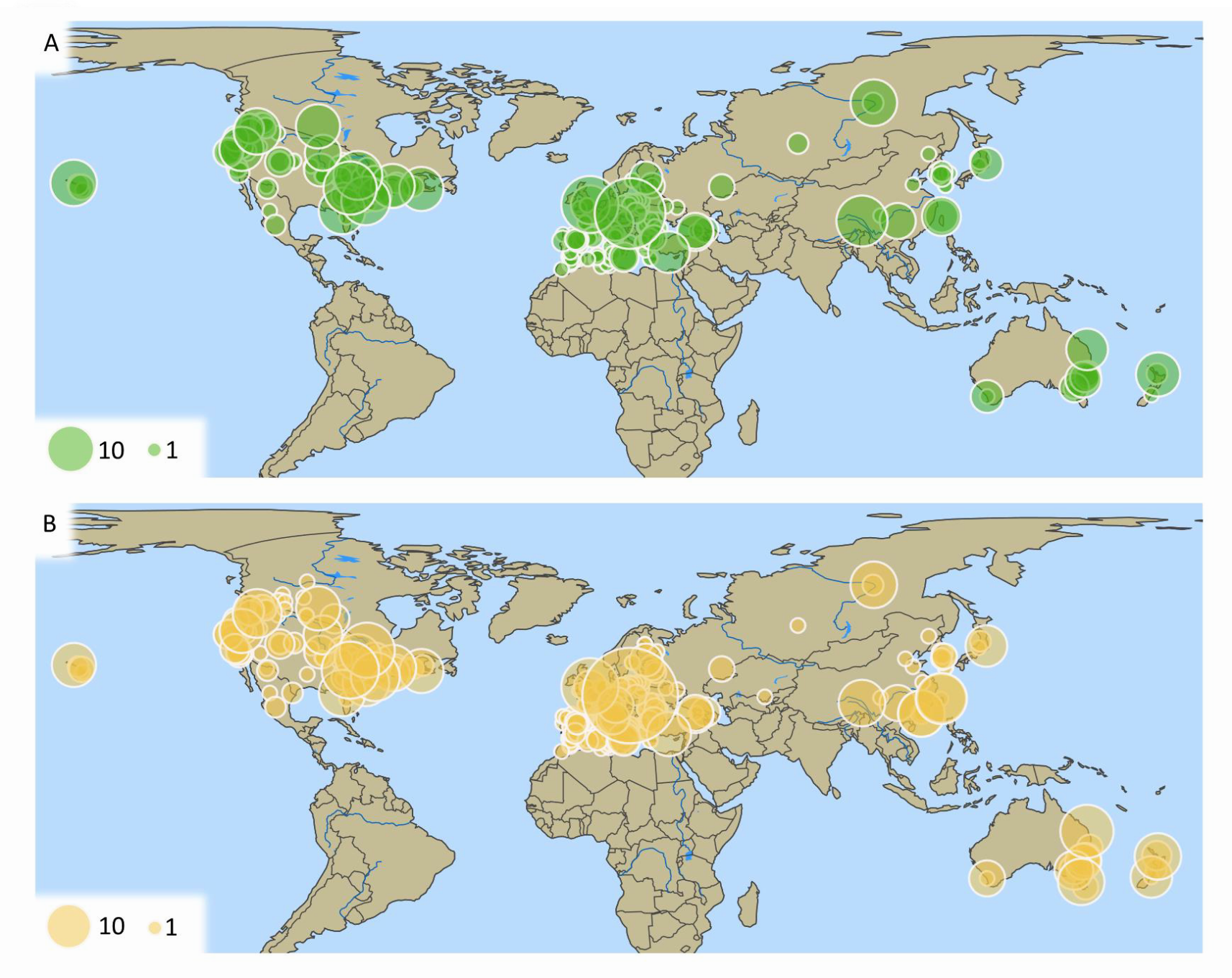
Sample size by location. **a**, ddRADseq (N = 559). **b**, mtDNA (N = 1,002). Size of points corresponds to sample size. Explore these data further through interactive data visualizations.

A total of 22,059 autosomal (ddRADseq) Single Nucleotide Polymorphisms (SNPs) for 559 individuals (average depth: 74X ± 28 sd; average missingness: 2.9% ± 4.3 sd) passed quality filtering (Fig 1a). We also sequenced a 502 bp region of the mitochondrial gene cytochrome c oxidase subunit 1 (*COI*) from 751 individuals (632 of these individuals were also used to generate ddRADseq data) and supplemented these sequences with 251 additional sequences from various online databases (total individuals with *COI* sequence = 1,002; Fig 1b).

### Global patterns of autosomal genetic differentiation and diversity

We filtered the ddRADseq data for autosomal markers and found evidence for at least seven genetically distinct clusters (ADMIXTURE lowest cross-validation error: 0.25 for K = 7) (Fig 2a). These genetic clusters largely correspond to the continental regions sampled and we refer to them henceforth as populations, named based on their sampling region: Europe, North Africa, Asia, Russia (east), North America (east), North America (west), Australia/New Zealand (Fig 2e). The greatest genetic differentiation was between Asia (including Russia (east)) and all other populations; average *F*_*ST*_ = 0.26±0.03sd (Fig 2c). Visual inspection of ancestry assignments (at higher values of K) suggests additional hierarchical levels of structure, primarily in Asia, but also within North America, and between Australia and New Zealand (Fig S2a,b). Surprisingly, we were unable to detect (geographically coherent) structure within Europe (except for Malta being distinct from the rest of Europe) or within Australia.

**Fig 2.**
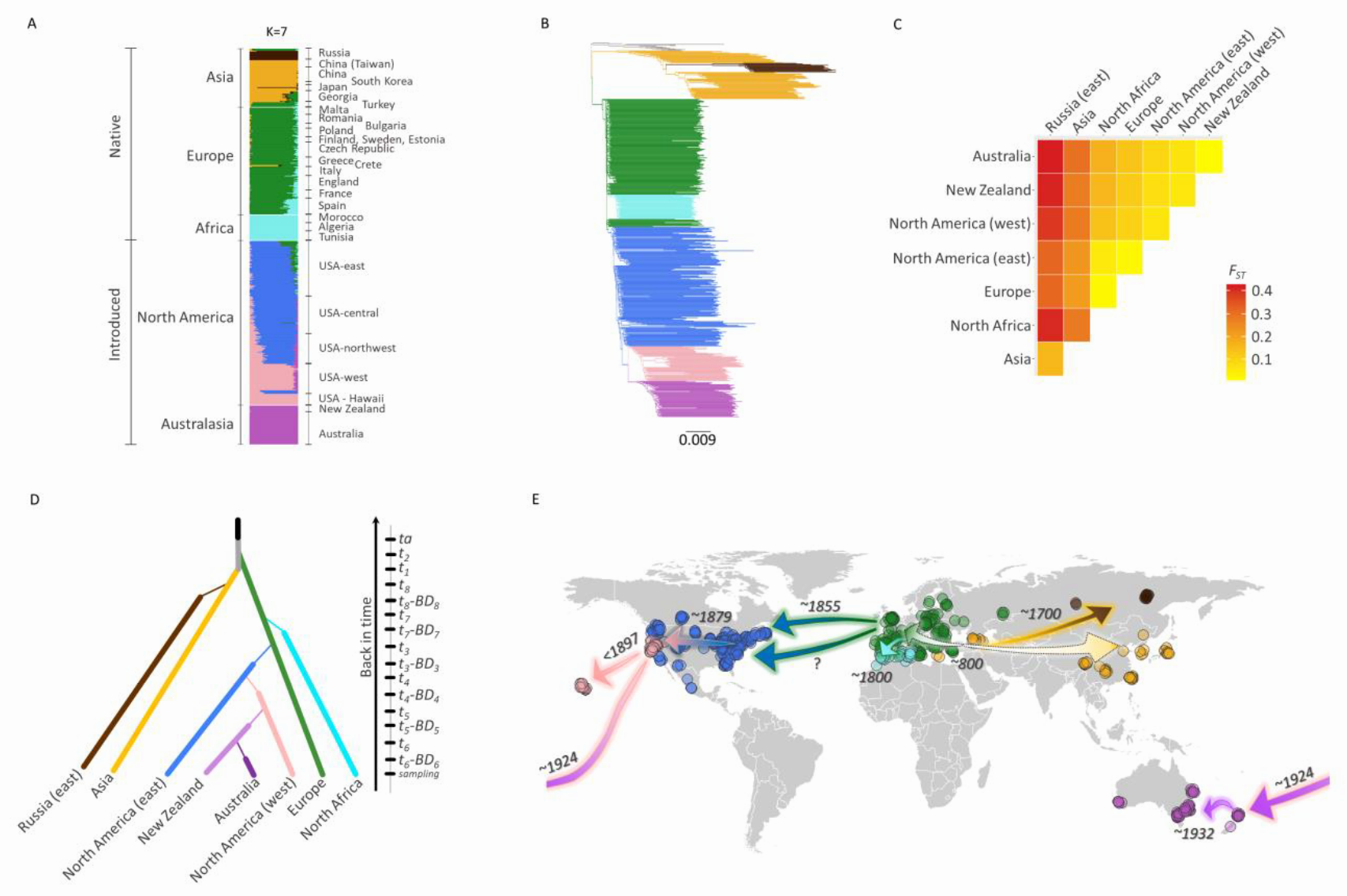
Global invasion history and patterns of genetic structure and diversity of *Pieris rapae*. **a**, Genetic ancestry assignments based on the program Admixture. **b**, Rooted neighbor-joining tree based on Nei’s genetic distance. **c**, Among population genetic differentiation based on Weir and Cockerham’s *F*_*ST*_; New Zealand and Australia are treated separately. **d**, Graphical illustration of divergence scenario chosen in ABC-RF analysis (Table 1), **e**, Geographic representation of divergence scenario with the highest likelihood based on ABC-RF analysis; points are colored based on their population assignment using Admixture (Fig 2a) and dates (Common Era) represent median estimates from ABC-RF analysis. All analysis based on 558 individuals genotyped for 17,917 ddRADseq SNPs. Explore these data further through <u>interactive data visualizations</u>.

Almost all recently introduced populations (i.e., North America, Australia and New Zealand) exhibit lower observed heterozygosity and nucleotide diversity compared with populations in the native range (i.e., Europe and Asia), consistent with population bottlenecks associated with these introductions (Fig 3). North America (east) was a notable exception among the introduced populations, with observed heterozygosity higher than populations found in the native range. All estimates of Tajima’s *D* fell within the range of −1 to 1, suggesting most populations are near equilibrium. However, there is a negative relationship between estimates of Tajima’s *D* and time since introduction—i.e., more recent introductions have higher (positive) estimates of Tajima’s *D*, suggesting that North America (west), New Zealand and Australia are still recovering from repeated population bottlenecks.

**Table 1.** Description of the competing scenarios and results of the six successive ABC analyses to infer the invasion history of *Pieris rapae*.

| Step | Scenario | Prior error rate |  |  | Random forest votes |  |  | Posterior probability |  |  |
| --- | --- | --- | --- | --- | --- | --- | --- | --- | --- | --- |
|  |  | Data 1 | Data 2 | Data 3 | Data 1 | Data 2 | Data 3 | Data 1 | Data 2 | Data 3 |
| *Analysis 1 - Native area - 18 summary statistics; 13,974 SNPs* |  | 13.82% | 14.49% | 14.29% |  |  |  |  |  |  |
|  | S1: Asia is the source of Europe |  |  |  | 57 | 188 | 207 | - | - | - |
|  | S2: Europe is the source of Asia |  |  |  | 132 | 132 | 43 | - | - | - |
|  | **S3: Asia and Europe both derived from an ancestral population** |  |  |  | **811** | **680** | **750** | **0.8479** | **0.8173** | **0.8353** |
| *Analysis 2 - Russia and North Africa - 115 summary statistics; 15,533 SNPs* |  | 16.26% | 17.22% | 17.07% |  |  |  |  |  |  |
|  | **S1: Asia and Europe are respectively the sources of Russia and Africa** |  |  |  | **602** | **676** | **521** | **0.7020** | **0.7549** | **0.6352** |
|  | S2: Asia and Africa are respectively the sources of Russia and Europe |  |  |  | 162 | 180 | 146 | - | - | - |
|  | S3: Africa and Russia are respectively the sources of Europe and Asia |  |  |  | 56 | 30 | 76 | - | - | - |
|  | S4: Europe and Russia are respectively the sources of Africa and Asia |  |  |  | 180 | 114 | 257 | - | - | - |
| *Analysis 3 - North America east (NAE) - 51 summary statistics; 16,753 SNPs* |  | 32.82% | 31.83% | 31.82% |  |  |  |  |  |  |
|  | S1: Asia is the source of NAE, 1 introduction |  |  |  | 0 | 9 | 2 | - | - | - |
|  | **S2: Europe is the source of NAE, 1 introduction** |  |  |  | **576** | **553** | **559** | **0.5010** | **0.6136** | **0.5064** |
|  | S3: Asia is the source of NAE, 2 introductions |  |  |  | 6 | 4 | 2 | - | - | - |
|  | S4: Europe is the source of NAE, 2 introductions |  |  |  | 418 | 434 | 437 | - | - | - |
| *Analysis 4 - North America west (NAW) - 116 summary statistics; 17,049 SNPs* |  | 11.44% | 11.54% | 10.95% |  |  |  |  |  |  |
|  | S1: Asia is the source of NAW |  |  |  | 19 | 35 | 13 | - | - | - |
|  | S2: Europe is the source of NAW |  |  |  | 144 | 121 | 41 | - | - | - |
|  | **S3: NAE is the source of NAW** |  |  |  | **721** | **720** | **933** | **0.8518** | **0.9288** | **0.9524** |
|  | S4: Europe is the source of NAW ~ 1600 CE |  |  |  | 85 | 73 | 7 | - | - | - |
|  | S5: Europe is the source of NAW ~ 1600 CE; NAW is the source of NAE |  |  |  | 31 | 51 | 6 | - | - | - |
| *Analysis 5 - New Zealand - 223 summary statistics; 17,100 SNPs* |  | 2.18% | 2.30% | 2.18% |  |  |  |  |  |  |
|  | S1: Asia is the source of New Zealand |  |  |  | 2 | 6 | 5 | - | - | - |
|  | S2: Europe is the source of New Zealand |  |  |  | 14 | 14 | 28 | - | - | - |
|  | S3: NAE is the source of New Zealand |  |  |  | 16 | 52 | 130 | - | - | - |
|  | **S4: NAW is the source of New Zealand** |  |  |  | **968** | **928** | **837** | **0.9739** | **0.9760** | **0.9802** |
| *Analysis 6 - Australia - 388 summary statistics; 17,116 SNPs* |  | 14.94% | 15.00% | 14.81% |  |  |  |  |  |  |
|  | **S1: New Zealand is the source of Australia** |  |  |  | **631** | **733** | **613** | **0.7797** | **0.8420** | **0.8124** |
|  | S2: NAW is the source of Australia |  |  |  | 63 | 33 | 62 | - | - | - |
|  | S3: New Zealand and Europe are the source of Australia (admixture) |  |  |  | 15 | 10 | 18 | - | - | - |
|  | S4: New Zealand and Asia are the source of Australia (admixture) |  |  |  | 15 | 7 | 10 | - | - | - |
|  | S5: New Zealand and NAW are the source of Australia (admixture) |  |  |  | 276 | 217 | 297 | - | - | - |

Results are provided for all three datasets. For each ABC analysis a forest of 1,000 trees was grown. The lines in bold characters corresponds to the selected (most likely) scenarios.

**Fig 3.**
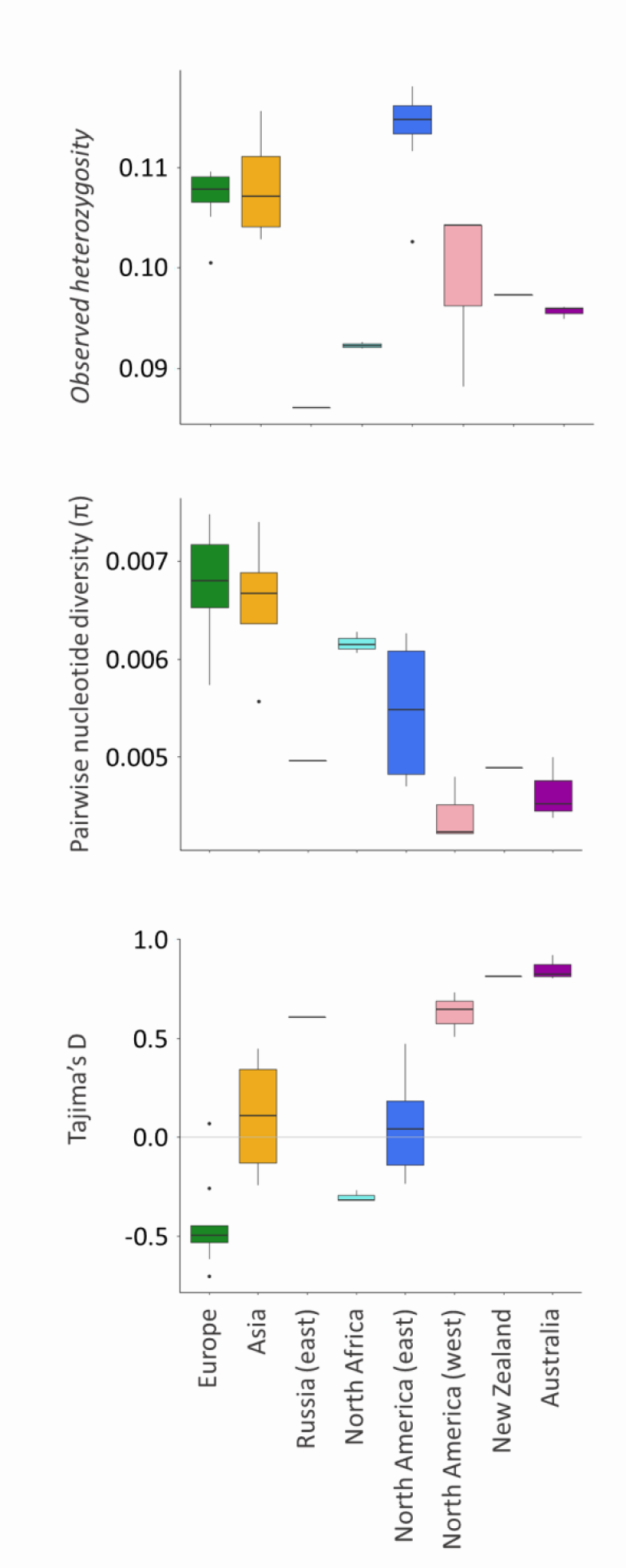
Patterns of autosomal genetic diversity—observed heterozygosity, pairwise nucleotide diversity, and Tajima’s *D*—by population.

### Global Invasion History

We compared a number of alternative invasion history scenarios for both the native and introduced populations using ddRADseq autosomal data within an approximate Bayesian computation random forest (ABC-RF) framework. We used an iterative process for selecting each bifurcation event, starting with native populations (Europe and Asia), then Russia (east) and North Africa, followed by the recently introduced populations (North America (east and west)), New Zealand and Australia, and then simulated a full model that incorporated all the best supported scenarios to get final parameter estimates (Fig 2e, Table 1). Based on this full final scenario (Fig 2d), posterior model checking revealed that the observed values of only six summary statistics out of 928 (i.e. 0.6%) fall in the tail of the probability distribution of statistics calculated from the posterior simulation (i.e. *p* < 0.05 or *p* > 0.95), which indicates that the chosen model ?tted well the observed genetic data. From parameter estimation (Table 1), we found the greatest support for a scenario with an ancestral population undergoing a demographic expansion *ca.* 20,000 (32,000–4,900) yrBP (Years Before Present) (Table S2). In evaluating the source for the Europe and Asia populations, we found the strongest support for the scenario of an ancestral population giving rise to both the Europe and Asia populations (∼85% posterior probability), *ca.* 1,200 (2,900–300) yrBP, over scenarios with Europe as the source for Asia, or Asia as the source for Europe (Fig S3a; Table 1). We evaluated multiple scenarios to determine the source for the Russia (east) and North Africa populations and found the strongest support for a scenario with Asia giving rise to the Russia (east) *ca.* 300 (800–200) yrBP, and the Europe population giving rise to the North Africa population ca. 200 (600–200) yrBP (Fig S3b; Table 1).

We found strong support (total of 996 random forest votes out of 1000) for Europe being the source of introduction to North America (east) (Fig S3c; Table 1). The scenario of a single introduction had only slightly better support than the scenario with multiple (two) introductions, and both have a similar number of random forest votes (576 and 418, respectively, out of 1,000, for dataset 1). Thus, we cannot clearly distinguish between these two scenarios, and prior error rate was consequently relatively high (∼33%). However, subsequent analyses performed by considering multiple introductions for the formation of North America (east) does not qualitatively change any of the following results (results not shown).

We found the strongest support for North America (east) serving as the source for the genetically distinct North America (west) population when compared to alternative scenarios with Asia or Europe (for both the scenario with ∼ 400 or 200 yrBP prior estimate for date of introduction) as the source (Fig S3d; Table 1). This introduction was estimated to have occurred *ca.* 137 yrBP. For the introduction into New Zealand, we found strong support for North America (west) being the source, when compared to Europe, Asia, or North America (east) as the source (Fig S3e; Table 1). The New Zealand population was found to have the greatest support as being the source for the introduction to Australia (Fig S3f; Table 1). All of these results were obtained with dataset 1 but were qualitatively confirmed by the analyses of datasets 2 and 3 (Table 1).

Demographic parameter estimates from ABC-RF analyses suggest each introduced population underwent a severe bottleneck, but the intensity (duration and number of founders with respect to the effective size of the source population) differed among populations (Table S2). Specifically, New Zealand and, more importantly, North America (west) were estimated to have undergone the most intense bottlenecks, whereas North America (east) and, to a lesser extent, Australia suffered less intense bottlenecks.

### Global patterns of mtDNA haplotype diversity and distribution

A total of 88 COI haplotypes were identified from 1,002 individuals, and 85% of these individuals harbored one of the eleven most common haplotypes (Fig 4; Fig S4, Table S3). The geographic distribution of mtDNA haplotypes is consistent with the invasion routes identified from ABC-RF analyses—haplotypes in introduced populations are largely a subset of those from putative source populations or differ by only one to two mutations from haplotypes in high frequency in the putative source populations (Fig 4a; see interactive figure to plot haplotypes individually— http://www.pierisproject.org/ResultsInvasionHistory.html).

**Fig 4.**
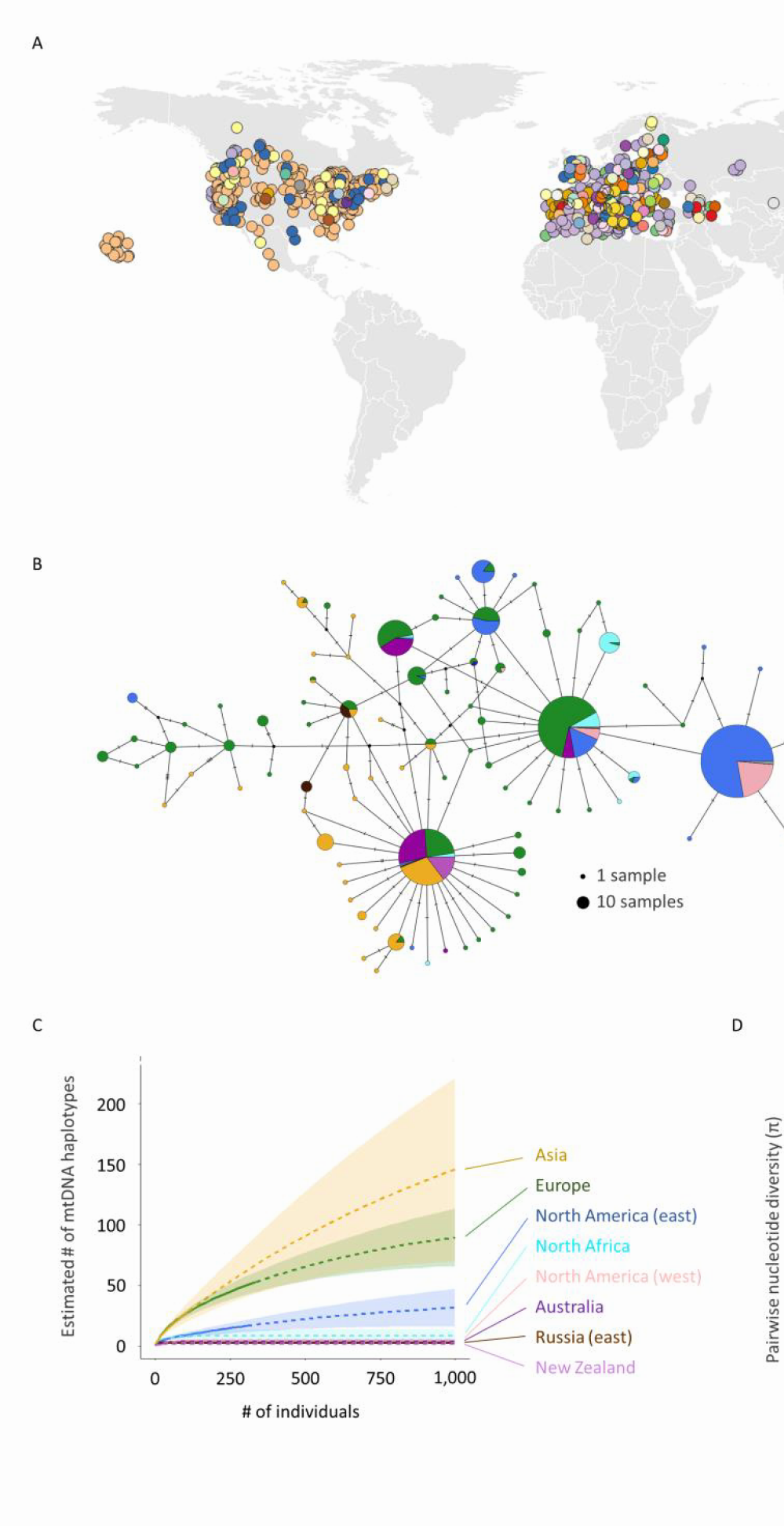
Global patterns of mitochondrial haplotype diversity. **a**, Geographic distribution of all 88 mtDNA haplotypes discovered (unique color for each haplotype; see <u>interactive data visualizations</u> to explore individual haplotypes); note points jittered to avoid overlapping (hidden) points, thus coordinates are approximate, and the color used for haplotypes are unrelated to those used in other panels. **b**, Haplotype network inferred using median-joining algorithm and colored by population. Hash marks between haplotypes represent base changes (mutations). **c**, Number of unique mtDNA haplotypes by population as well as subpopulation estimated using a rarefaction approach (see Methods) and plotted by geographic location. **d**, Pairwise nucleotide diversity by population. Explore these data further through <u>interactive data visualizations</u>.

Estimates of mtDNA haplotype diversity (richness) were highest in Asia and Europe and had large confidence intervals (based on rarefaction curve analysis), indicating these populations were likely undersampled (Fig 4c). All introduced populations had substantially lower estimates of mtDNA haplotype diversity. New Zealand, Australia, North America (west), North Africa, and Russia (east) were estimated to have less than a dozen mtDNA haplotypes, whereas North America (east) had substantially (∼3X) more mtDNA haplotypes and was significantly greater than all other introduced populations (based on non-overlapping confidence intervals). Estimates of mtDNA nucleotide diversity are similar to those observed for autosomal markers using ddRADseq data, with the exception of Australia, which had higher nucleotide diversity than New Zealand and North America (west) (Fig 4d). From a global perspective, there appears to be a general trend of decreasing nucleotide diversity with increasing distance from southern Europe and the eastern Mediterranean region (Fig S5).

Considering the mtDNA haplotypes found in North America and their frequencies in sub-populations of Europe, we estimate that the minimum number of individuals that would need to have been sampled from the sub-population of England (2.3) is 23 ± 12 sd individuals or 123 ± 88 sd individuals for Spain/southern France (2.4) to account for all haplotypes in North America.

## Discussion

### Citizen science greatly expands range-wide sampling of butterflies

We show for the first time how the public can contribute to our understanding of species invasions through a collection-based citizen science project. Collections made by citizen scientists dramatically expanded the geographic scope of our study by nearly doubling the number of countries we were able to include in our analyses. Samples from many of these countries either solely or primarily came from citizen scientists, including nearly the entire region of Australasia. Moreover, these contributions allowed us to increase substantially the total number of individuals in each of the populations studied, with nearly half of all specimens sequenced in our study coming from citizen scientists. We estimate that the use of citizen science to aid in the collection of *P. rapae* from across its near-global range resulted in tens of thousands of dollars (USD) in cost-savings that would have been required to cover salary and travel costs. We believe our citizen science approach can be applied to other systems, particularly to organisms that are easily identifiable (e.g., spotted lantern fly or Giant African snail) and easy to transport (e.g., dead invertebrates), to address questions in invasion biology as well as a broad range of questions in ecology and evolutionary biology. As examples, we currently are leveraging our large collection to address questions concerning the effects of climate and land use changes on wing pigmentation of this butterfly and to identify genomic regions underlying ecological selection.

The development, implementation, and maintenance of this project was not trivial, as is the case with many citizen science projects, and required considerable time and effort engaging the public (e.g., contacting organizations, using social media, responding to emails) and processing samples. Collections-based citizen science projects that focus on less charismatic species or incorporate non-lethal forms of sampling (e.g., eDNA) and are easy to collect (slow moving or sessile) may have the greatest success. We suggest that those interested in applying a collections-based citizen science approach seek advice from, or build collaborations with, individuals with experience in the field of citizen science.

### Geographic spread and divergence of <u>P. rapae</u> driven by host plant domestication

The Europe and Asia populations of *P. rapae* are believed to represent distinct subspecies—*P. rapae rapae* and *P. rapae crucivora*, respectively—based on phenotypic differences^9^ and evidence for reproductive isolation^15^. Our study provides further support for this, revealing the two main genetic lineages recovered by ddRADseq for *P. rapae* worldwide correspond to the Europe (*Pieris rapae rapae*) and Asia (*Pieris rapae crucivora*) populations. It has long been hypothesized that the domestication of brassicaceous crops, which are the primary host plants for this butterfly, aided the spread and/or divergence of the Europe and Asia subspecies^10^. Our data support this hypothesis. We estimate the divergence of both the Europe and Asia populations (subspecies) occurring within the last ∼3000 years. This estimate overlaps with estimates for the domestication of brassicaceous crops that occurred during this time period (i.e., *Brassica oleracea* and *Brassica rapa*)^16–18^. However, our results do not support the hypothesis that Europe was the source of the Asia population^9^. Instead, our results suggest both the Europe and Asia populations independently diverged from an ancestral population, most likely *ca.* 1,200 yrBP. This time period corresponds with the intensification in the cultivation of *B. oleracea* varieties, such as cabbage and brussels sprouts^19^.

It remains unclear whether *P. rapae* spread across and occupied Europe and Asia during this expansion event, and then diverged *in situ* in response to the domestication of brassicaceous crops, or whether *P. rapae* was more restricted in distribution (e.g., confined to Europe or the eastern Mediterranean region) and diverged in association with the domestication of brassicaceous crops across Eurasia. Consistent with the hypothesis that Europe and Asia *P. rapae* populations diverged as they spread out of the eastern Mediterranean, the range boundaries of the Europe and Asia populations (subspecies) abut in the eastern Mediterranean region and genetic diversity generally decreases with increasing distance from this region. In further support of this hypothesis, there is growing evidence that the domestication of *Brassica oleracea* and *Brassica rapa* originated in the Mediterranean region^18^. However, additional sampling in western Asia is needed to further evaluate this hypothesis.

Interestingly, the putative ancestral population that gave rise to the Europe and Asia populations appears to have undergone a rapid increase in effective population size *ca.* 7,000– 28,000 yrBP. This time period overlaps with early human development of agriculture. However, our median estimate for the date of this expansion was *ca.* 20,000 yrBP, placing it at the end of the last glacial maximum *ca.* 23,000–19,000 yrBP^20^. Changes in the distribution and demography of species in response to glacial–interglacial cycles is well documented^21,22^, and may be more likely to have facilitated a major demographic shift in *P. rapae*, as the earliest domestication of brassicaceous crops was relatively recent (earliest evidence being *ca.* 7,000 yrBP)^17^.

### Recent invasion history largely reflects historical records, but with a few unexpected findings

Although historical records of species invasions can be misleading^23^, our molecular genomics-based reconstruction of the *P. rapae* global invasion history is largely consistent with that historically documented through observations. As expected, we found Europe to be the most likely source of this butterfly’s introduction into North America. However, we unexpectedly found that there was no discernable nuclear genetic structure in Europe (even when K = 30), making it impossible to narrow down with confidence the source population to a specific locality or country (e.g., England vs Spain). However, mtDNA haplotype distributions and frequencies in European countries suggest England as the most likely source—i.e., fewer individuals would be required to produce the mtDNA haplotypes found in North America, if England was the source rather than Spain and southern France. We do not know what specific factors account for the lack of genetic structure in Europe. One possibility is long-distance dispersal of this species^24^ coupled with historic and/or ongoing human assisted dispersal has led to high levels of gene flow. Another interesting possibility supported by some evidence^24–26^ is that *P. rapae* is migratory or undergoes migratory-like events.

Historical records obtained by Samuel Scudder pointed to possible multiple introductions of *P. rapae* into North America, which occurred during or shortly after its initial invasion^11^. Confirming multiple introductions from the same source population early in an invasion, particularly one that quickly underwent a rapid expansion, is extremely difficult. The best-fit model to our data suggests a scenario with a single introduction, but it seems reasonable there were multiple introductions for a couple of reasons. First, both competing scenarios—one vs multiple introductions from Europe—had a similar number of random forest votes (576 vs 418 out of 1,000) and the selected scenario (i.e. one introduction from Europe) had a low posterior probability estimate (∼0.5; in contrast, all scenarios that were chosen in the other analyses had posterior probability estimates >0.70). Second, we estimated a rather low bottleneck intensity, with a founding population size of ∼50-100 individuals; this estimate is much higher than a previous estimate of one to four individuals^27^. It seems unlikely that North America was founded from a single introduction given this rather large estimated founding population size and the reasonable assumption that no more than a few dozen (unrelated) butterflies would be transported on any one ship. Third, multiple introduction events (from Europe) would help explain the higher heterozygosity found in North America (east) than in the native range—i.e., multiple introductions aided in the rebound in genetic diversity as the introduced population spread across eastern North America.

Our second unexpected finding was the identification of a genetically distinct population within North America that is restricted to the western USA. We found evidence of admixture in areas where the two North America (east and west) populations come into contact, suggesting that these genetically distinct populations are neither geographically or reproductively isolated from each other. The geographic extent of this admixture zone is not clear from our sampling, nor are the consequences of gene flow between these populations. We initially hypothesized this western population represents an early introduction brought by Spaniards during the 1600s, but our data instead indicate that it most likely results from a secondary founder event from the North America (east) population brought during the ∼1860-1880s as a result of the rapid development of railroad lines^28^, specifically from the eastern USA to central California (Video S1).

Our results confirm previous speculation that North America (west), likely San Francisco, California, was the source of introduction to the Hawaii islands, based on individuals from Hawaii being assigned to the North America (west) cluster but not being reported in the Hawaiian islands until 1987^29^ (after the arrival of *P. rapae* to central California). Also, unexpectedly, our results suggest that the introduction of *P. rapae* to New Zealand came from North America (west), and not from Europe, as was believed to be the most likely case given the United Kingdom was the largest exporter into New Zealand at the time^30^. Lastly, previous speculation that New Zealand is the immediate source of *P. rapae* in Australia^13^ is supported by our data.

### Implications for invasion biology

Our study also sheds light more broadly on invasion biology. Growing evidence shows many invasive populations are able to flourish and adapt to new environments despite substantial loss of genetic diversity—a phenomenon termed the genetic paradox of invasions^31^. *Pieris rapae* is a remarkable example of this paradox. This butterfly has rapidly expanded its range following each new founding event, despite repeated population bottlenecks—at least four separate times, with each new founding population the product of a previously bottlenecked population (i.e., multiple serial founding events). Whether introduced populations maintained high genetic variation in ecologically relevant traits following each founding event remains unclear. Evidence of local adaption for thermal tolerances among populations in North America^32^ suggests such variation exists. However, resolving this paradox and the persistent puzzle of how this butterfly has been an extremely successful invader into new environments will require future studies to assess the relative contributions of factors such as adaptative evolution, phenotypic plasticity, natural enemy escape, and domestication of its host plants.

## Methods

### Specimens collection and DNA extractions

The *Pieris rapae* specimens were collected as part of an international citizen science project—Pieris Project (pierisproject.org)—that was launched in June 2014 and through collections by researchers. A website was created in 2014 for Pieris Project that included a description of the research goals and collection protocol—specimens were to be individually placed in hand-made or glassine envelopes, labeled with location and date collected, and placed in a freezer overnight, then air-dried for at least two days and shipped using standard mail. The project was advertised through social media—Twitter (@PierisProject) and Facebook (https://www.facebook.com/pierisproject/), and through listservs, social media, and blogs of Entomological and Lepidopterists societies and nature/science/citizen science related organizations (e.g., YourWildLife, eButterfly, National Geographic, and SciStarter). Once received, specimens were stored in 95% ethanol and kept at −20 °C; depending on the collector, specimens were air-dried for a few days to years prior to being placed in ethanol. Genomic DNA was isolated from tissue from prothorax or (2-3) legs using DNeasy Blood and Tissue Kit spin-columns (Qiagen, Hilden; Cat No./ID: 6950).

To estimate the contributions by scientists, we binned the collector of each specimen into one of two categories: 1) researcher, and 2) citizen scientist. Collectors whose identity was known (>90% of participants) were not considered as citizen scientists if they had a college degree in biology. This makes our estimated contribution by citizen scientists a conservative estimate, as some of these participants may consider themselves citizen scientists. There is a great deal of debate as to what does or does not constitute being a citizen scientist. Our threshold is based on our previously stated definition of citizen science that is accepted by many within the field of citizen science.

### ddRADseq sequencing and filtering

Nine reduced-complexity libraries were generated using a double-digest restriction-site-associated DNA fragment procedure (ddRAD)^33^ following Ryan et al., 2018^34^. Briefly, genomic DNA (∼400 ng) was digested with the restriction enzymes EcoR1 and Mse1 and a universal Mse1 and barcoded EcoR1 adapter ligated to the digested DNA. Ligated products were diluted 10 times with 0.1X TE buffer prior to PCR enrichment. Amplified products with unique barcodes were pooled into a single mixture prior to purification. The library was purified three times with 0.8X volume of Agencourt Ampure XP beads (Beckman Coulter, A63881). At the end of each round of purification, elution volume was reduced to 0.25 – 0.5X of the beginning sample volume. After three rounds of purification, each library (1.0 µg) was size selected for 400 to 600 bp fragment length using 1.5% DF Cassette and BluePippin System (Sage Science). Libraries were evaluated by Bioanalyzer 2100 system and sequenced across one lane Illumina MiSeq (University of Notre Dame, Genomics Core Facility), 14 lanes of Illumina HiSeq 4000 (12 at University of Illinois and 2 at Beijing Genomics Institute); most samples were sequenced on 2 (some 3) independent lanes.

Raw reads were demultiplexed and barcodes/cutsite removed using a custom Python script. Reads were further trimmed and cleaned with the program Trimmomatic (v0.32)^35^ using default settings. The first 5 bp and any after 80 bps were then trimmed from all reads and only reads at least 76 bp in length were retained, resulting in all reads being exactly 76 bp.

Reads were then aligned to the *Pieris rapae* genome v1 ^27^ using BWA-aln (v0.7.15)^36^. Variant calling was performed using GATK’s Haplotypecaller (v3.8)^37,38^ with the default settings. Filters were applied in the following order: kept only biallelic SNPs, applied GATK’s “hard filtering” (QD < 2.0 ‖ MQ < 40.0 ‖ MQRankSum < −12.5 ‖ ReadPosRankSum < −8.0), SNPs with a genotype quality (GQ) < 20 were converted to missing data, removed SNPs with minor allele frequency less than 0.01, kept SNPs with min of 1X coverage for 50% of individuals, removed SNPs with coverage > 95^th^ percentile (112.8 X coverage), removed individuals with > 75% missing data, kept only SNPs with a minimum of 10X coverage in 90% of individuals, and removed individuals with >25% missing data. Finally, SNPs with heterozygosity > 0.6 were considered potential paralogs and were discarded.

As there is no linkage map for *P. rapae* and the genome is not assembled into chromosomes, we applied a simple heterozygosity method to determine whether SNPs were autosomal or sex (Z) linked. To do this we used the expectation that females should be homozygous at all SNPs on the Z-chromosome—females are the heterogametic sex (ZW) in Lepidoptera. Using 231 females, we calculated the percentage that were heterozygous or homozygous at each site (SNP) using the is_het function from the R package vcfR (v1.6.0)^39^ and custom scripts in R. SNPs with greater than 25% missing data were removed. A scaffold (and all SNPs within) was considered putatively Z-linked if > 60% of the SNPs fell below the threshold of having less than 1% of the females being heterozygous (average number of SNPs for each scaffold was 84 ± 101; mode = 4).

To complement the heterozygosity method, we also inferred the chromosome assignment of each *P. rapae* scaffold using the approach by Ryan et al. (2017)^40^. Briefly, we used blastx (ncbi-blast-2.2.30+)^41,42^ to blast all peptide sequences within each scaffold of the *P. rapae* assembly against the *Bombyx mori* genome (silkdb v2.0)^43^. The *B. mori* scaffold with the most significant blast hits (based on Bit scores) was retained and used to determine the putative chromosome of each *P. rapae* scaffold. All, except one scaffold, of those we identified as Z-linked using the heterozygosity method, mapped to chromosome 1 (Z) and 2 (W) of *B. mori* (Fig S6). That we found some regions of *P. rapae* mapping to the *B. mori* chromosome 2 (W) suggests they are not completely syntenic—the *P. rapae* genome was assembled from males and thus there should be no scaffold mapping to this chromosome. Some *P. rapae* scaffolds mapping to chromosome 2 (W) of *B. mori* were recovered as actually being on the chromosome 1 (Z) based on the heterozygosity method.

Using a subset of the putative Z-linked markers—SNPs where < 1% of females had a heterozygous call (i.e., SNPs with a high likelihood of being Z-linked)—we validated the sex of each individual. Females and males with >20% or <20% of these SNPs being heterozygous were considered possibly mislabeled males and females respectively. These individuals were flagged, and the specimens double-checked visually; in all cases, visual identification confirmed that these individuals were mislabeled.

### Inference of Population Structure and Diversity

Population structure was investigated with the model-based clustering algorithm ADMIXTURE^44^ using default settings and a cross-validation = 10 for K 1-30 using a modified SNP dataset, i.e., pruned for LD (*r*^*2*^ > 0.2; calculated using VCFtools geno-r2) (17,917 SNPs). The optimal K was that with the lowest cross validation error. The Bayesian program fastSTRUCTURE^45^ as well as a non-model-based multivariate approach—Discriminant Analysis of Principal Components (DAPC; Fig S2c)^46^—were also used to confirm the results from ADMIXTURE (see Supplementary Materials for more details) using the R package adegenet (v2.1.1)^47^. Genetic assignments were plotted using custom scripts and the R package pophelper (v2.2.3)^48^. A Neighbor-Joining tree based on genetic distance was constructed in the poppr (v2.8.0)^49^ and ape (v5.1)^50^ packages in R, that included the species *Pieris napi, Pieris brassicae*, and *Pieris canidia* as outgroups, using only sites with at least 15X coverage in 90% individuals from this new dataset. Trees were visualized using FigTree (v1.4.4)^51^. Population differentiation was estimated between all populations using the Weir & Cockerham’s estimator of *F* ^52^ implemented in VCFtools (v0.1.15) using 10 kb windows and a window step size of 5 kb.

All measures of genetic diversity (*H*_*obs*_, π, *and* Tajima’s *D*) were calculated using SNPs restricted to scaffolds longer than 100kb (22,059 SNPs). In an attempt to minimize the Wahlund effect (i.e., reduction of heterozygosity caused by subpopulation structure), individuals were split into spatially contiguous subpopulations from within the seven identified by ADMIXTURE (N=34; one subpopulation from Mexico was not included because it contained only three individuals); these were the same subpopulations used for the ABC-RF analyses. To control for differences in sample size, we computed each statistic 1,000 times using a random subset (without replacement) of seven individuals (size of smallest population). Heterozygosity was estimated using the R package adegenet v2.1.1. Calculations for *π, and* Tajima’s *D* were estimated using a dataset containing invariant sites (i.e., vcf files were created using gatk-4.0.4.0 with the flag -allSites true and the same filters as described above were then applied) with VCFtools (v0.1.15) using 10 kb windows (and a window step size of 5 kb used for estimating *π*).

### ABC-RF-based inferences of global invasion history

An approximate Bayesian computation analysis (ABC)^53^ was carried out to infer the global invasion history of *Pieris rapae*. The eight populations considered in the ABC analysis corresponded to the seven identified by ADMIXTURE, with an additional separation of New Zealand and Australia for geographical reasons. Each population was represented in the analysis by a single sub-population (individuals sampled within the same subregion and within a three-year period) (dataset 1). ABC is a model-based Bayesian method allowing posterior probabilities of historical scenarios to be computed, based on historical data and massive simulations of genetic data. We used historical information (e.g., dates of first observation of invasive populations) to define 6 sets of competing introduction scenarios that were analyzed sequentially (Table 1 and Fig S3). Step by step, subsequent analyses used the results obtained from the previous analyses, until the most recent invasive populations were considered. The first set of competing scenarios (three scenarios) considered the evolutionary relationship between the Asian and European populations. In the second analysis (four scenarios), we explored the links between Asia, Europe, North Africa and Russia (east). In the third analysis (four scenarios), we set North America (east) as the target and determined whether it originated from Asia or Europe, either through one or two introductions. In the fourth analysis (5 scenarios), North America (west) could be originating either from Europe, Asia or North America (east), and the introduction could be ancient (400 yrBP) in the case of Europe. In the fifth analysis (four scenarios), New Zealand could be originating either from Europe, Asia, North America (east) or North America (west). Finally, the sixth analysis (five scenarios) aimed at deciphering the origin of the Australian population by testing as source population New Zealand, North America (west), and admixtures between New Zealand and either Europe, Asia or North America (west). All scenarios of all analyses are detailed in Table 1 and Fig S3.

In our ABC analysis, historical and demographic parameter values for simulations were drawn from prior distributions defined from historical data and demographic parameter values available from empirical studies on *Pieris rapae*^11–13,29^, as described in Table S5. Simulated and observed datasets were summarized using the whole set of summary statistics proposed by DIYABC^54^ for SNP markers, describing genetic variation per population (e.g., mean gene diversity across loci), per pair (e.g., mean across loci of *F*_ST_ distances), or per triplet (e.g., mean across loci of admixture estimates) of populations (see the DIYABC v2.1.0 for details about statistics), plus the linear discriminant analysis axes^55^ as additional summary statistics (Table S6). The total number of summary statistics ranged from 18 to 388 depending on the analysis (Table 1).

To compare the scenarios, we used a random forest process^56,57^. Random forest is a machine-learning algorithm which uses hundreds of bootstrapped decision trees to perform classification using a set of predictor variables, here the summary statistics. Some simulations are not used in tree building at each bootstrap (i.e., the out-of-bag simulations) and can thus be used to compute the “prior error rate”, which provides a direct method for cross-validation. We simulated a 10,000 SNPs datasets for each competing scenario using the standard Hudson’s algorithm for minor allele frequency (i.e., only polymorphic SNPs over the entire dataset are considered), so the number of used markers ranged between 13,974 and 17,116 depending on the analysis (Table 1). We then grew a classification forest of 1,000 trees based on the simulated datasets. The random forest computation applied to the observed dataset provides a classification vote (i.e., the number of times a model is selected among the 1,000 decision trees). The scenario with the highest classification vote was selected as the most likely scenario, and we then estimated its posterior probability by way of a second random forest procedure of 1,000 trees^57^. To evaluate the global performance of our ABC-RF scenario choice, we computed the prior error rate based on the available out-of-bag simulations and conducted the complete scenario selection analysis with two additional datasets with different sub-populations (dataset 2 and dataset 3) representative of the same populations as dataset 1^58^.

We then performed a posterior model checking analysis on a full final scenario including all 8 populations (dataset 1), to determine whether this scenario matches well with the observed genetic data. Brie?y, if a model ?ts the observed data correctly, then data simulated under this model with parameters drawn from their posterior distribution should be close to the observed data. The lack of ?t of the model to the data with respect to the posterior predictive distribution can be measured by determining the frequency at which the observed summary statistics are extreme with respect to the simulated summary statistics distribution (hence de?ning a tail-area probability or *p*-value, for each summary statistic). We simulated 100,000 data sets under the full ?nal scenario (17,609 SNP and 928 summary statistics), and then obtained a ‘posterior sample’ of 5,000 values of the posterior distributions of parameters through a rejection step based on Euclidean distances and a linear regression post-treatment^53^. We simulated 1,000 new datasets with parameter values drawn from this “posterior sample”, and each observed summary statistic was compared with the distribution of the 1,000 simulated test statistics, and its *p*-value, corrected for multiple comparisons with the false discovery rate procedure^59^, was computed.

Finally, 10,000 simulated datasets of the full final scenario were used to infer posterior distribution values of all parameters, and some relevant composite parameters under a regression by random forest methodology^60^, with classification forests of 1,000 trees. The simulation steps, the computation of summary statistics, as well as the model checking analysis were performed using DIYABC v2.1.0. All scenario comparisons and parameter estimations were carried out in R using the package abcrf (v1.7.1)^57^.

### mtDNA sequencing and analysis

A 1,600 bp region of COI was amplified using primers optimized to work with multiple species within the genera *Pieris* (Pieridae_COI_F 5-AAATTTACAATYTATCGCTTA-3, Pieridae_COI_R 5-TGGGGTTTAAATCCATTACATATW-3). When these primers failed we amplified a 658 bp region of COI using previously published primers^61^. PCR amplicons were purified using magnetic beads and amplified using standard fluorescent cycle sequencing PCR reactions (ABI Prism Big Dye terminator chemistry, Applied Biosystems). Sequencing reactions were purified using Agencourt CleanSeq magnetic beads (Beckman Coulter) and run on an ABI-3730XL-96 capillary sequencer (Applied Biosystems) at the University of Florida biotechnology facility (ICBR) or Macrogen (Macrogen Inc). Individuals with both forward and reverse reads were assembled in Geneious 11.0.4 using the De Novo Assemble tool with default settings. The find heterozygotes tool (peak similarity set to 50%) was used to find and discard any sequences found to be heterozygous. Reads were trimmed to 502 bp and aligned (error probability limit of 0.001) with sequences from GenBank and Barcode of Life databases using MUSCLE Alignment in Geneious with default settings.

To evaluate whether we were adequately sampling mtDNA haplotype diversity, we plotted rarefaction curves (estimates of haplotype richness by sampling effort) for each population using iNEXT^62^ and predicted the total haplotypes for each population assuming 1,000 sampled individuals. A median-joining haplotype network was created using POPART^63^ for all populations and for each population separately.

In an effort to further pinpoint whether the introductions in North America came from western (i.e., United Kingdom) or southwestern (i.e., Spain and France) Europe, we estimated the minimum number of individuals that would need to be sampled from each of these native populations to generate the mtDNA diversity found in North America. Specifically, for each native subpopulation, we randomly sampled (with replacement) a haplotype from each subpopulation based on their haplotype frequencies, until all haplotypes represented in North America were sampled and simulated this procedure 10,000 times for each subpopulation. This approach assumes that the true source population will be the most parsimonious—i.e., require sampling of fewer individuals to create the diversity found in North America.

## Acknowledgements

We would like to thank all the participants in the Pieris Project, without their help this research would not have been possible. We would also like to thank Arthur Shapiro for his extraordinary insights into this system and to the many researchers who contributed specimens (see full list here). A special thank you to Sang-guy Park for donating specimens from his private collection. Many thanks to Jacqueline Lopez and Melissa Stephens in the Notre Dame Genomics & Bioinformatics Core Facility for ddRADseq library preparations. This research was funded by a USDA-NIFA Post Doctoral Fellowships grant #2017-67012-26999 to S.F.R. A.E. is supported by the NSF Graduate Research Fellowship Program (GRFP). R.V. was supported by project CGL2016-76322-P (AEI/FEDER, UE). G.T. is supported by the MINECO programme IJCI-2016-29083 and by the National Geographic Society (grant WW1-300R-18). E.A.H. was supported by a Marie Curie Actions IO Fellowship no. 330136.

## Data Availability

Demultiplexed ddRADseq reads generated in this study are available through NCBI’s Sequence Read Archive associated with Bioproject (<ID>, SRA: <ID>). All new *COI* sequences were deposited to the Barcode of Life Database (BOLD; <ID>). All metadata and scripts associated with analyses in this study have been deposited on DRYAD (<link>).

## Contributions

Author contributions: S.F.R. designed the research project. S.F.R conceived of, created, implemented, and runs the citizen science project—Pieris Project; S.F.R. performed all molecular work, with assistance from M.M.D. with prepping ddRAD libraries and M.A.R. with developing mtDNA primers; S.F.R. performed genomic diversity and structure analyses; E.L. conducted ABC-RF analyses with assistance from S.F.R.; S.F.R., R.V., G.T., V.D., A.E., E.A.H. contributed specimens and/or mtDNA sequences; S.F.R., M.W.E. and A.E. designed educational material related to the citizen science project; and S.F.R. wrote the manuscript with contributions from E.L., A.E., R.V., G.T., V.D., M.A.R., M.M.D., M.W.E., E.A.H., Y.Y., M.E.P., and D.D.S. All authors read and approved the final manuscript.

## Supplementary Tables

**Table S1.**
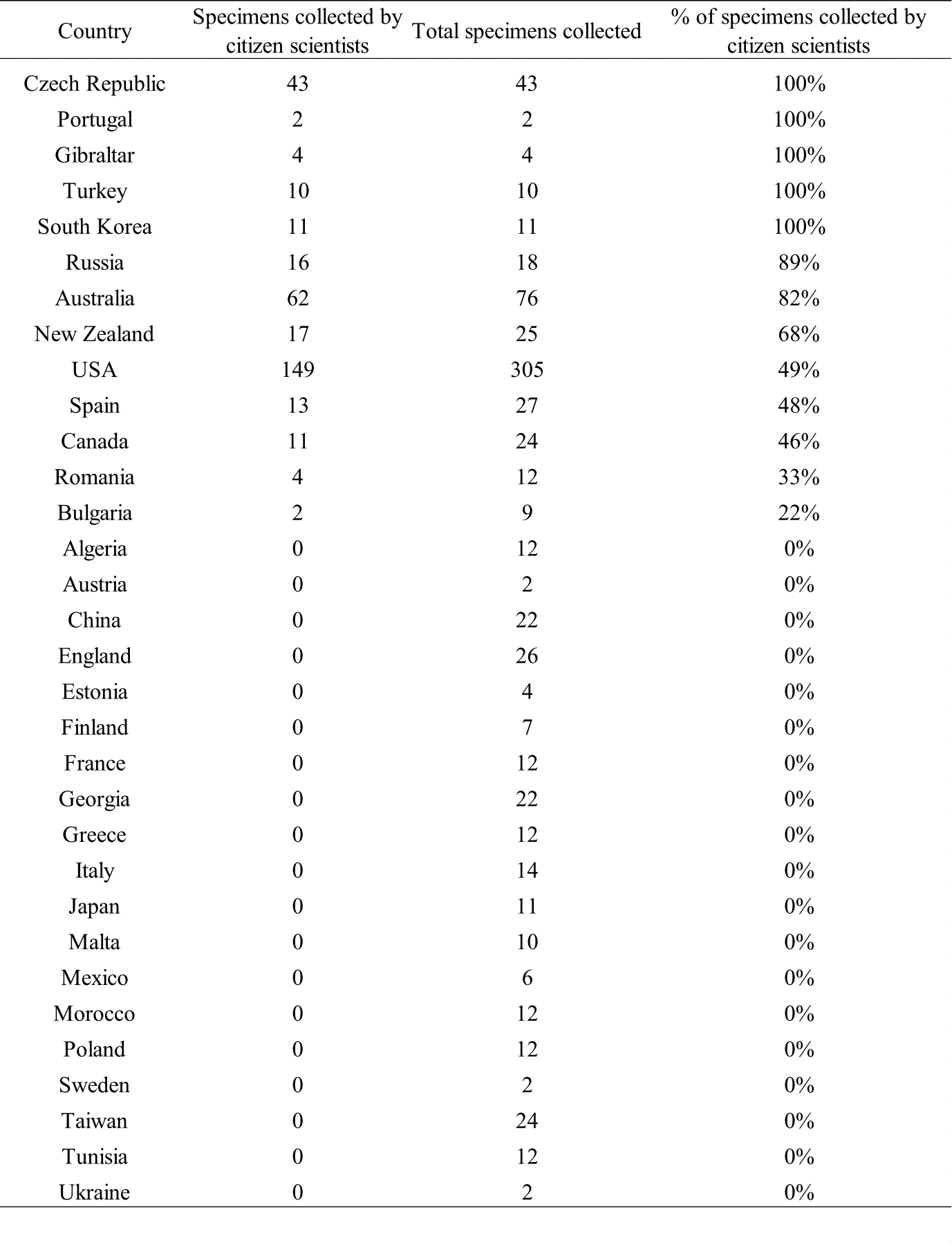
Contribution of specimens made by citizen scientists.

**Table S2.**
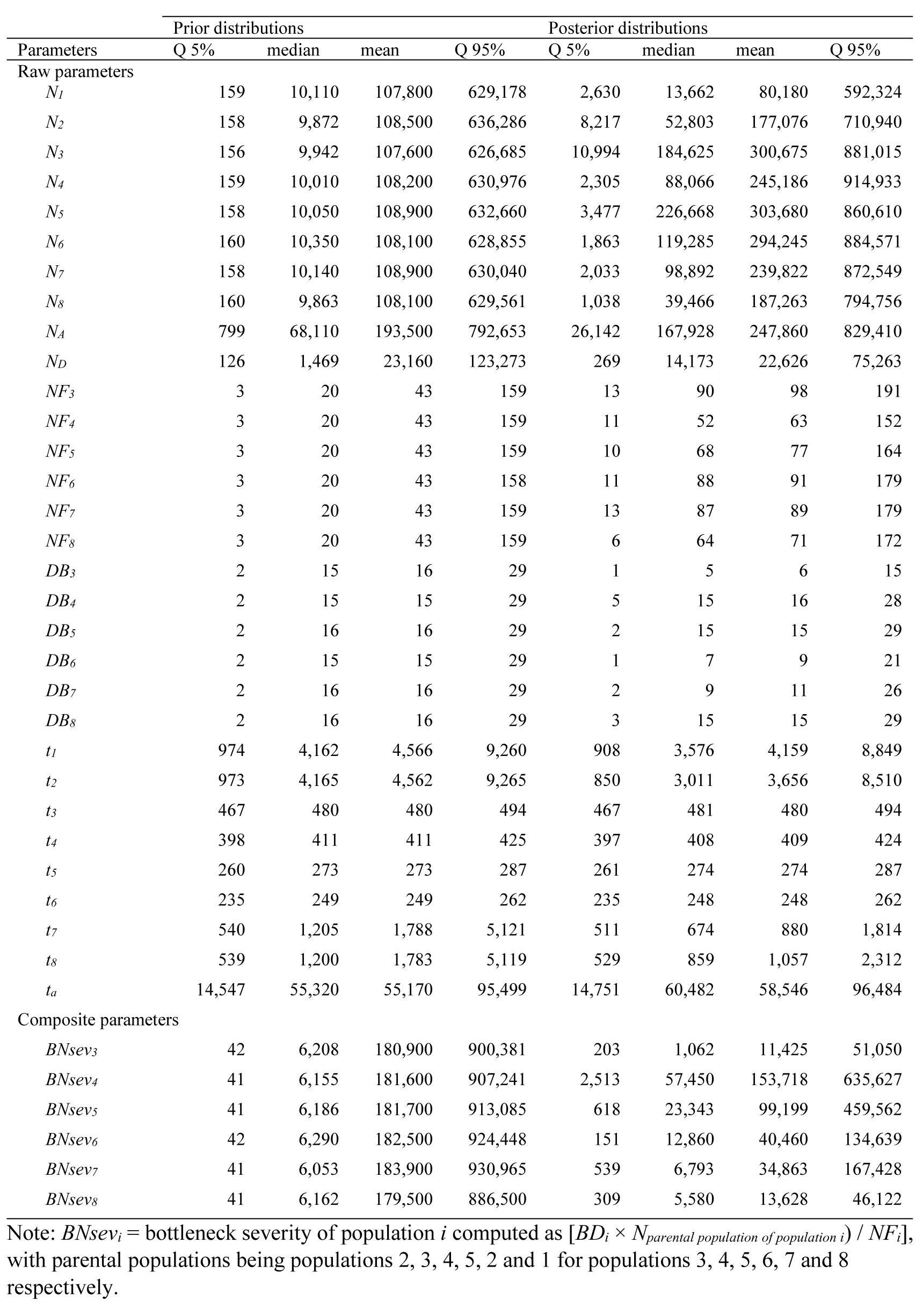
Prior and posterior distributions of all parameters and several composite parameters of the full final complete scenario (Fig 2d) performed with dataset 1.

**Table S3.** Metadata for specimens used in this study.

Included as supplementary file (too large)

**Table S4.** Populations used for ABC-RF analyses.

Included as supplementary file (too large)

**Table S5.**
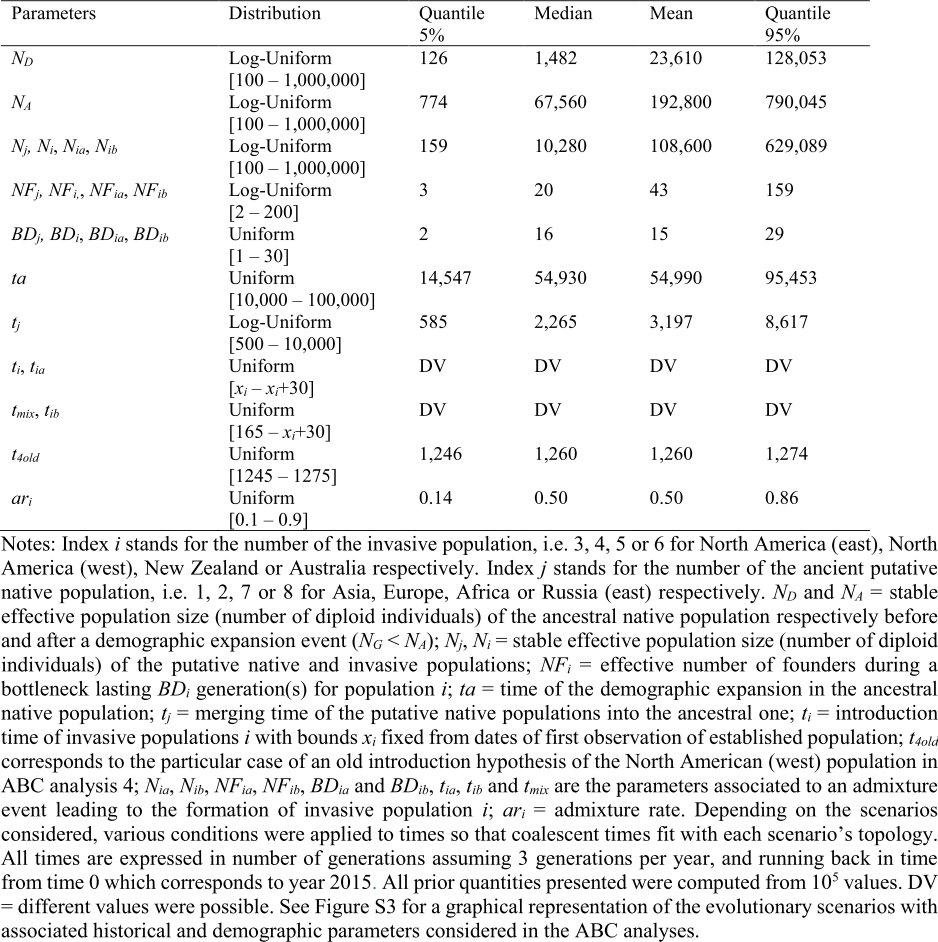
Prior distributions of demographic and historical parameters used in ABC analyses processed to retrace the worldwide invasion routes of Pieris rapae.

**Table S6.**
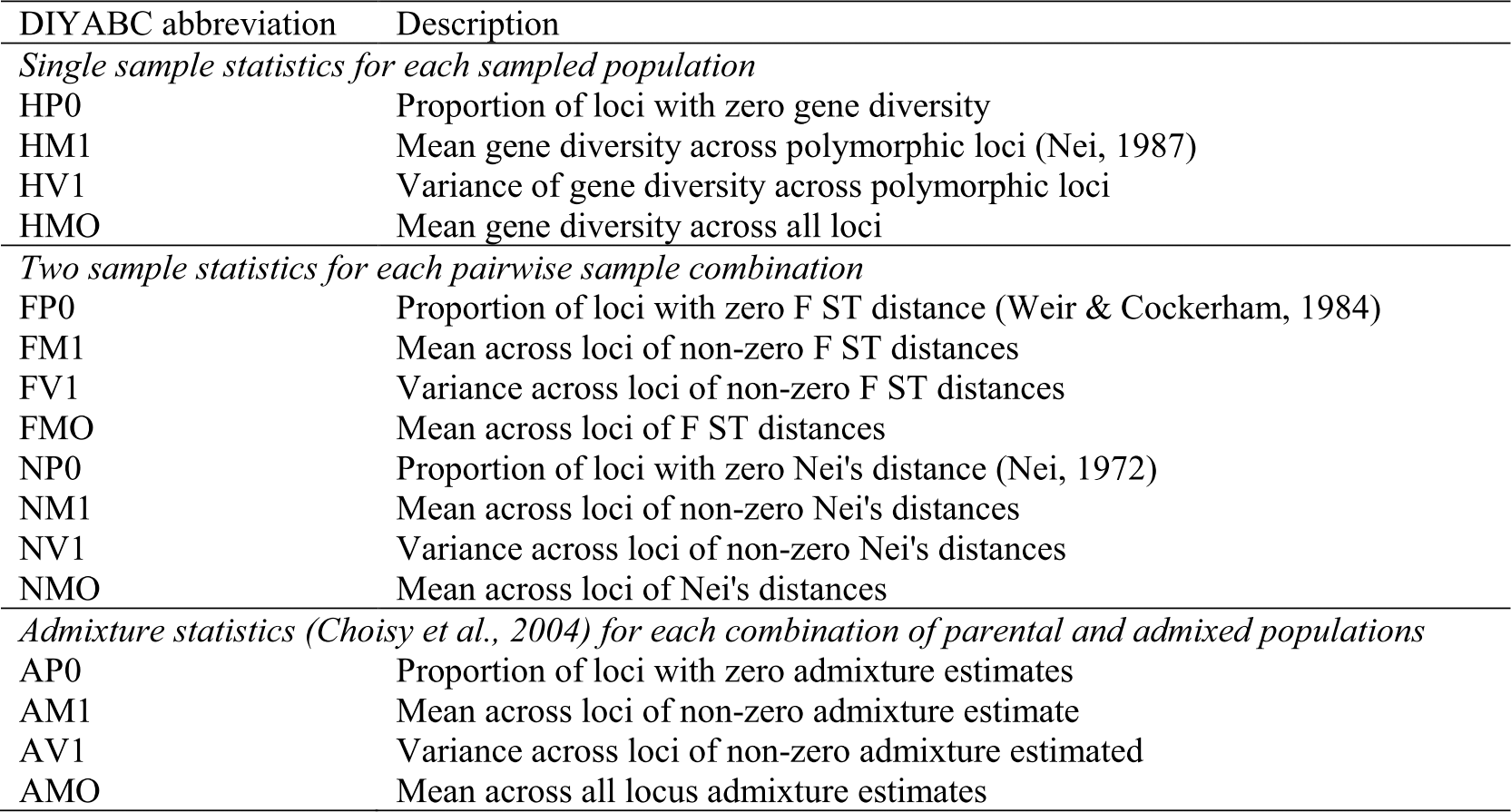
Summary statistics used in all DIYABC simulations (Cornuet et al., 2014).

## Supplementary Figures

**Fig S1.**
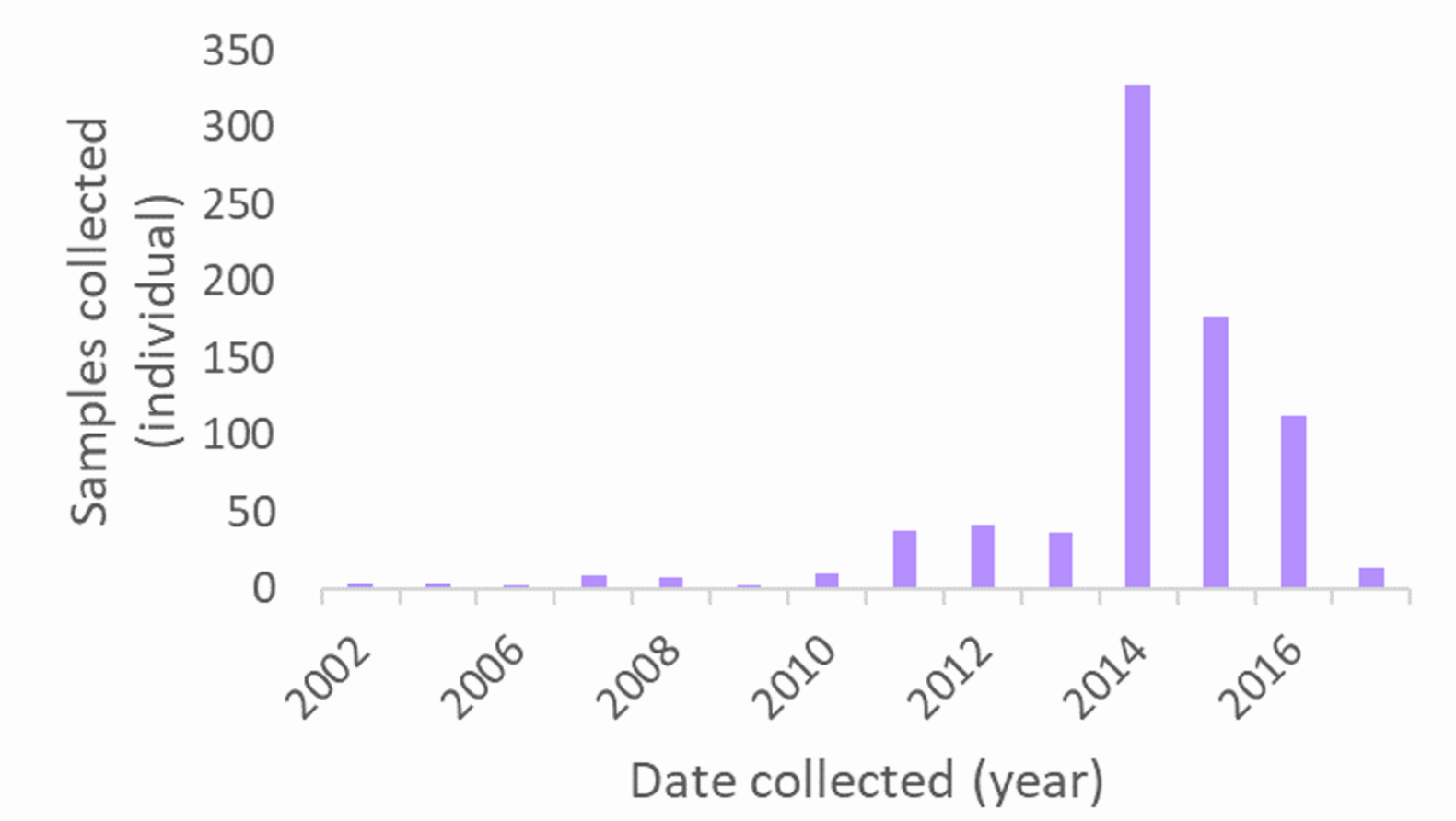
Sample sizes of *Pieris rapae* specimens by year collected.

**Fig S2.**
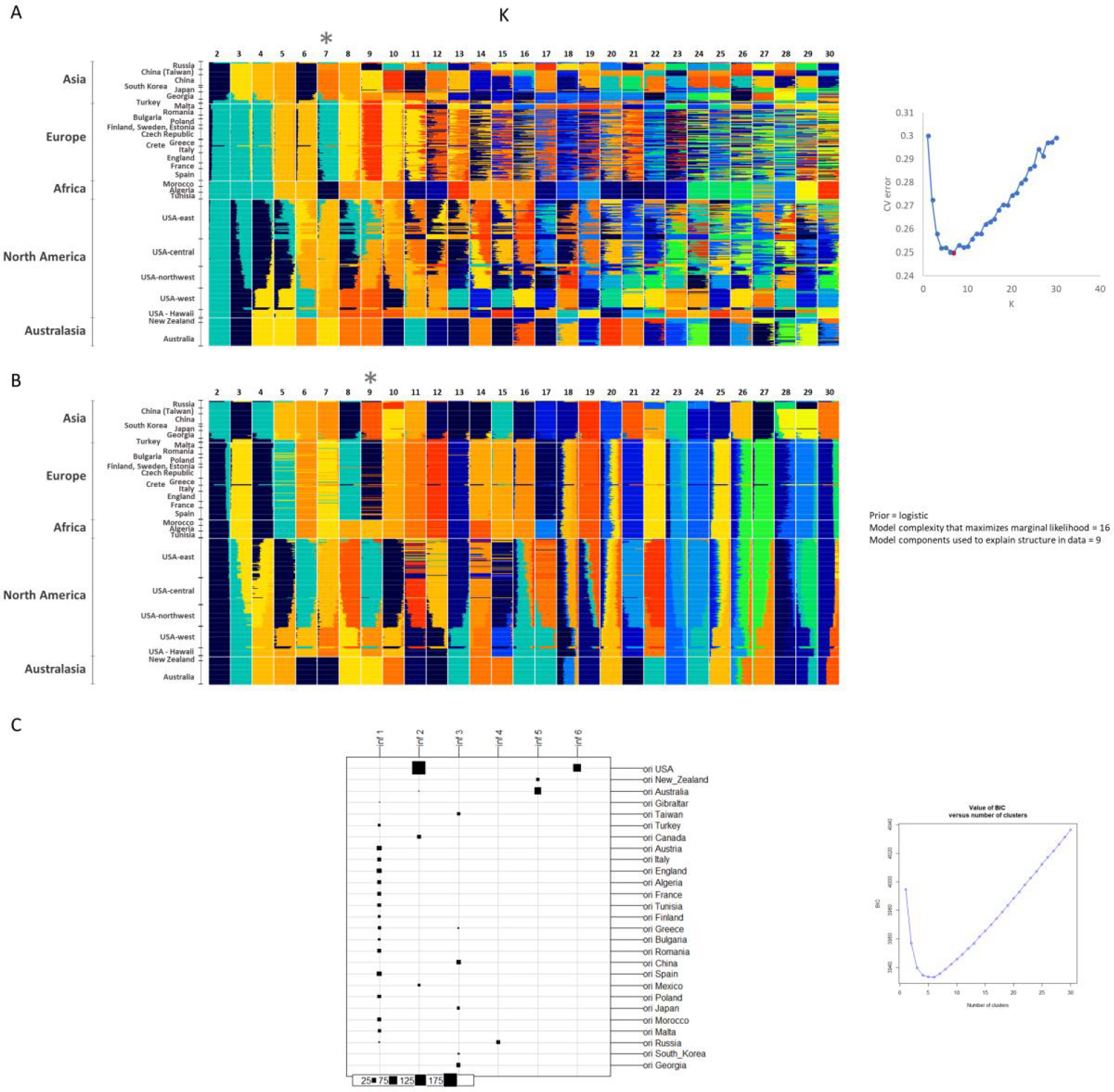
Population ancestry assignment plots for K:2-30, using **a**, ADMIXTURE, **b**, fastSTRUCTURE, and **c**, Discriminant Analysis of Principal Components (DAPC). For each analysis the evaluation for optimal K is included.

**Fig S3.**
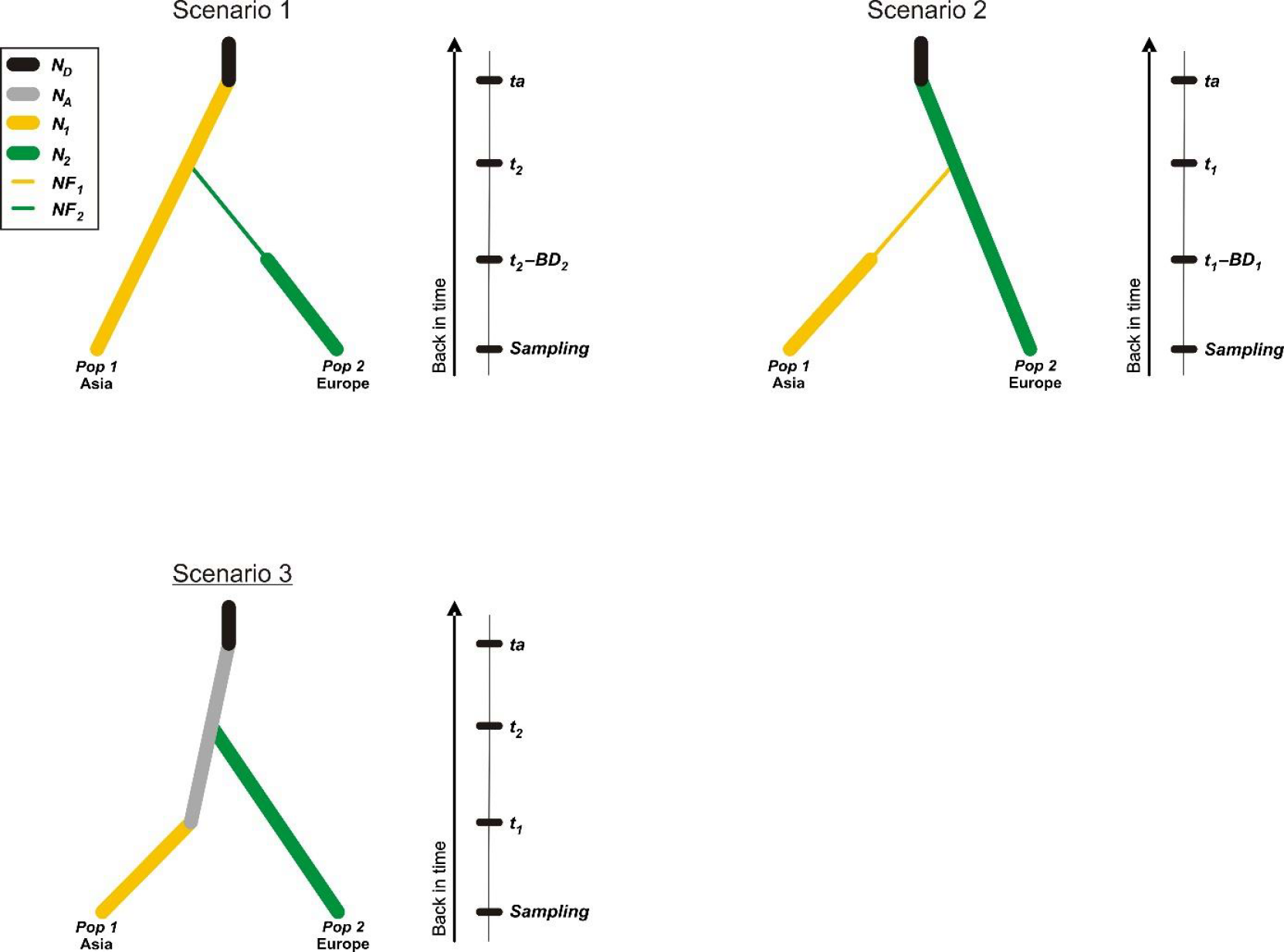
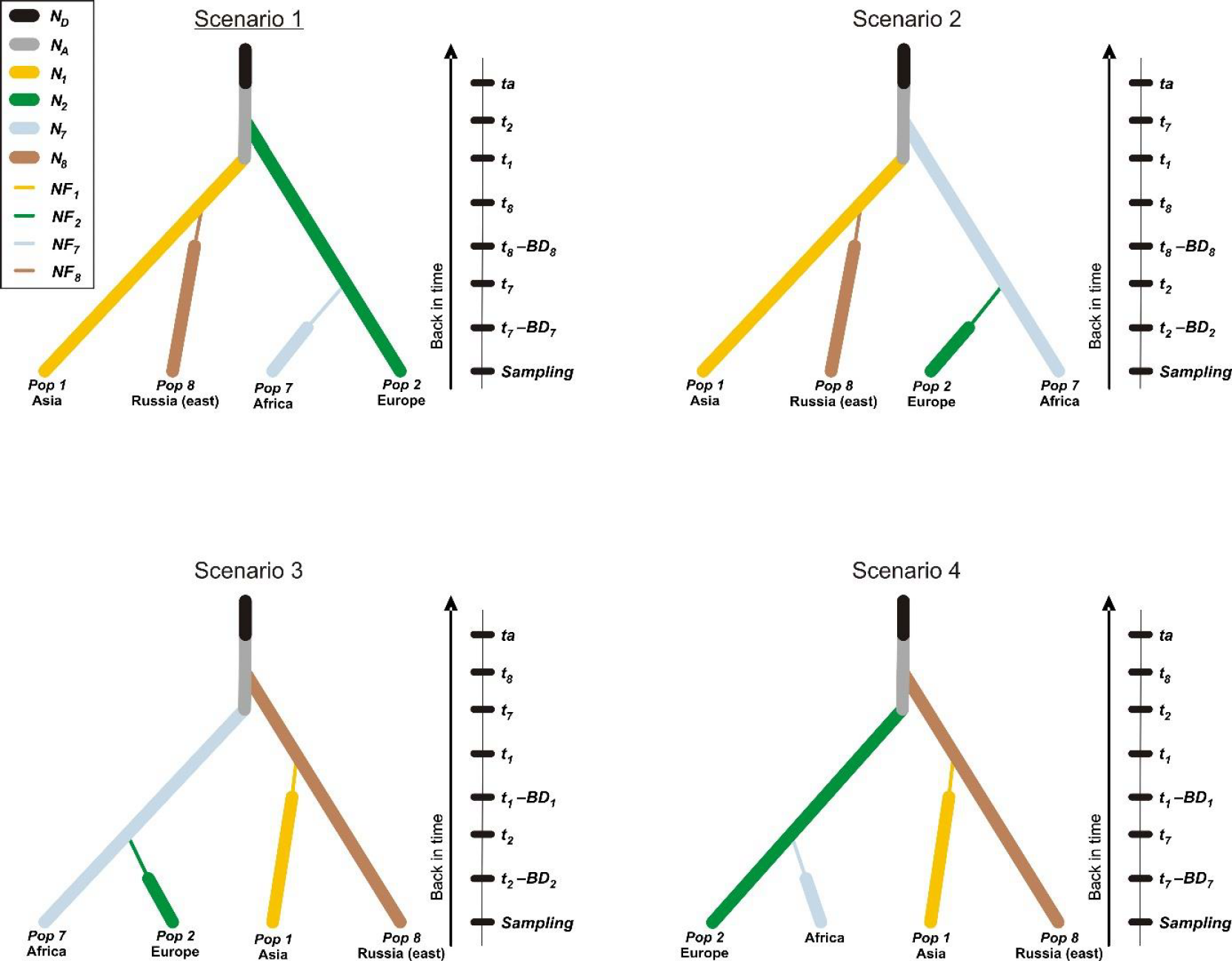
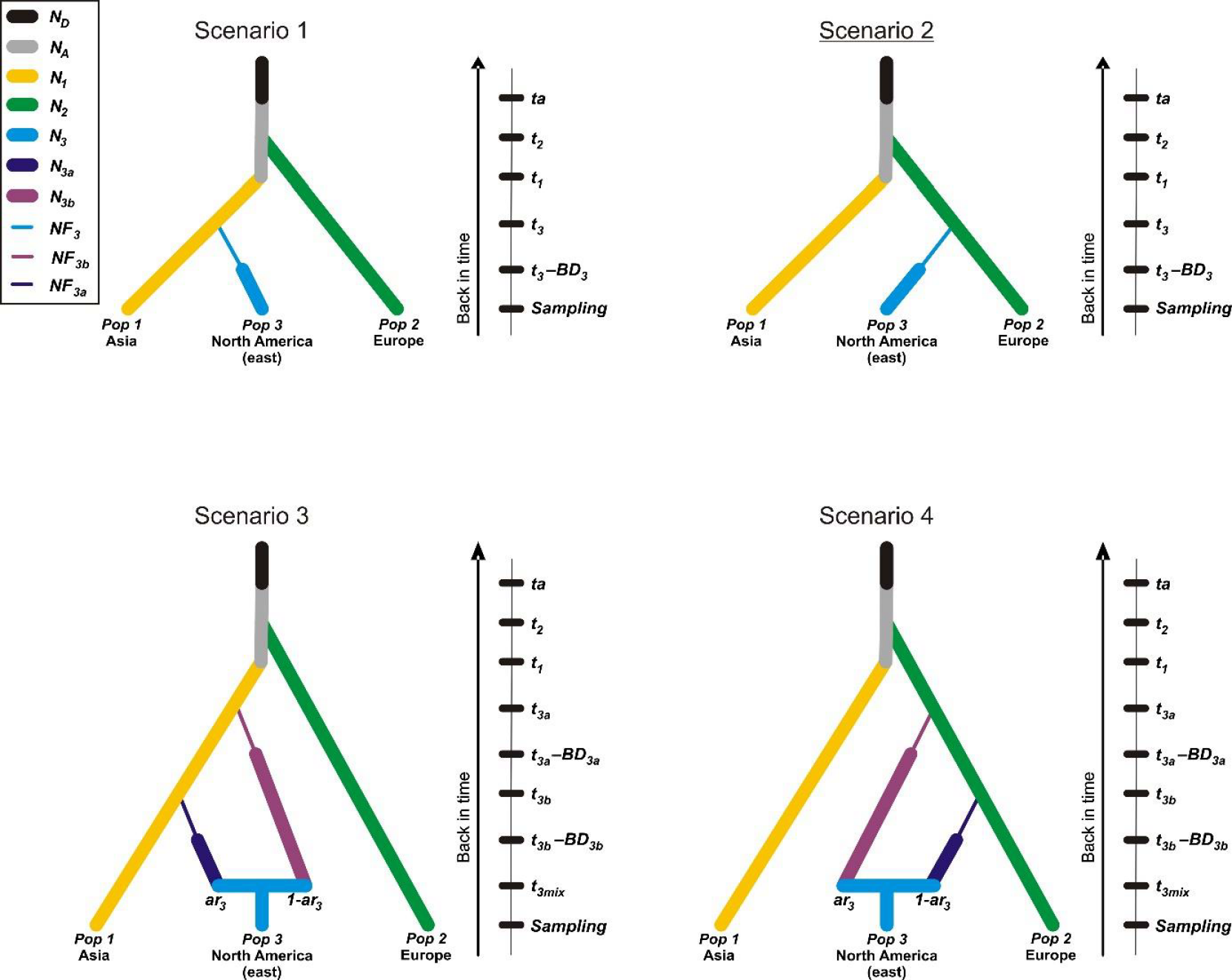
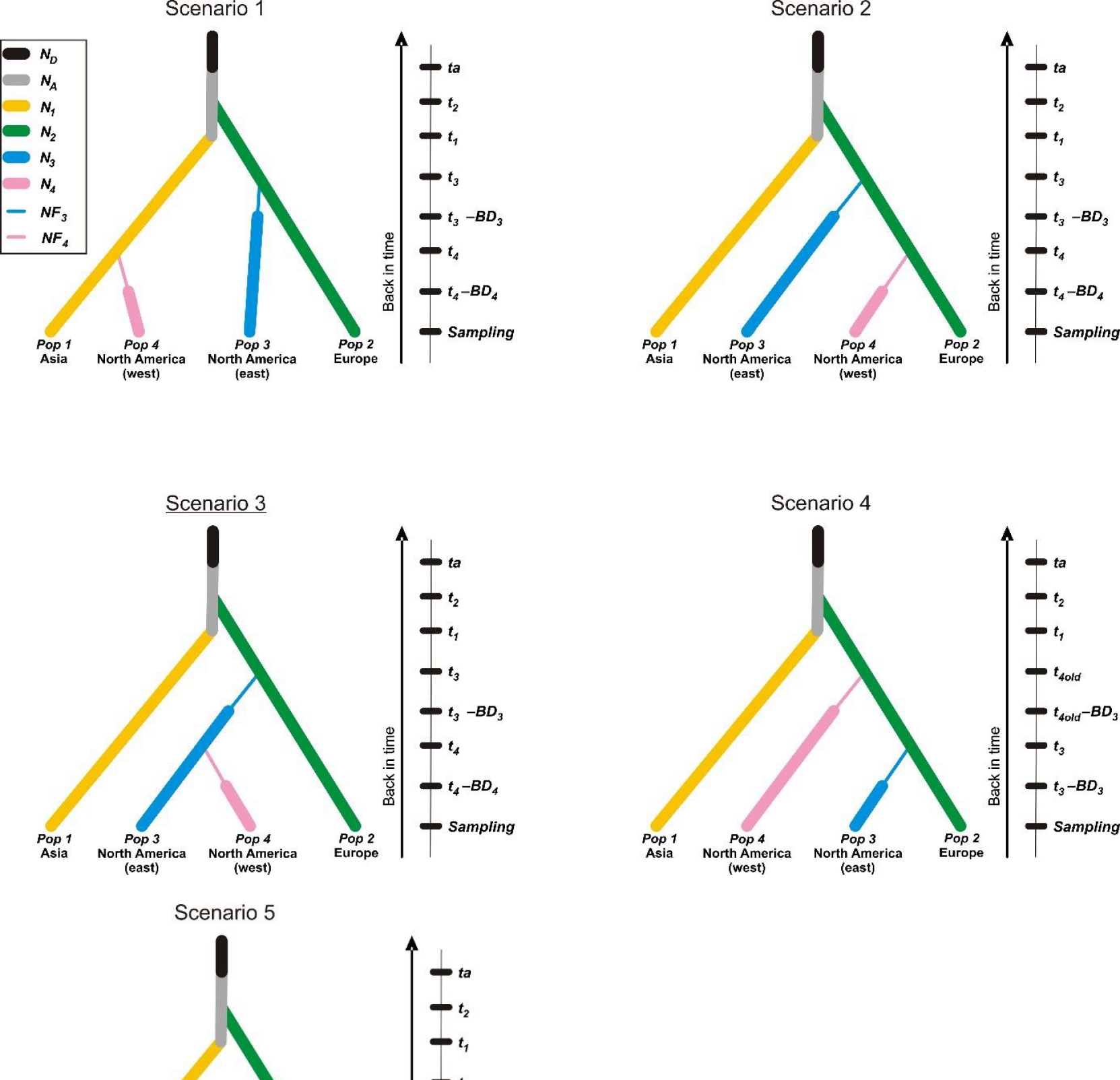
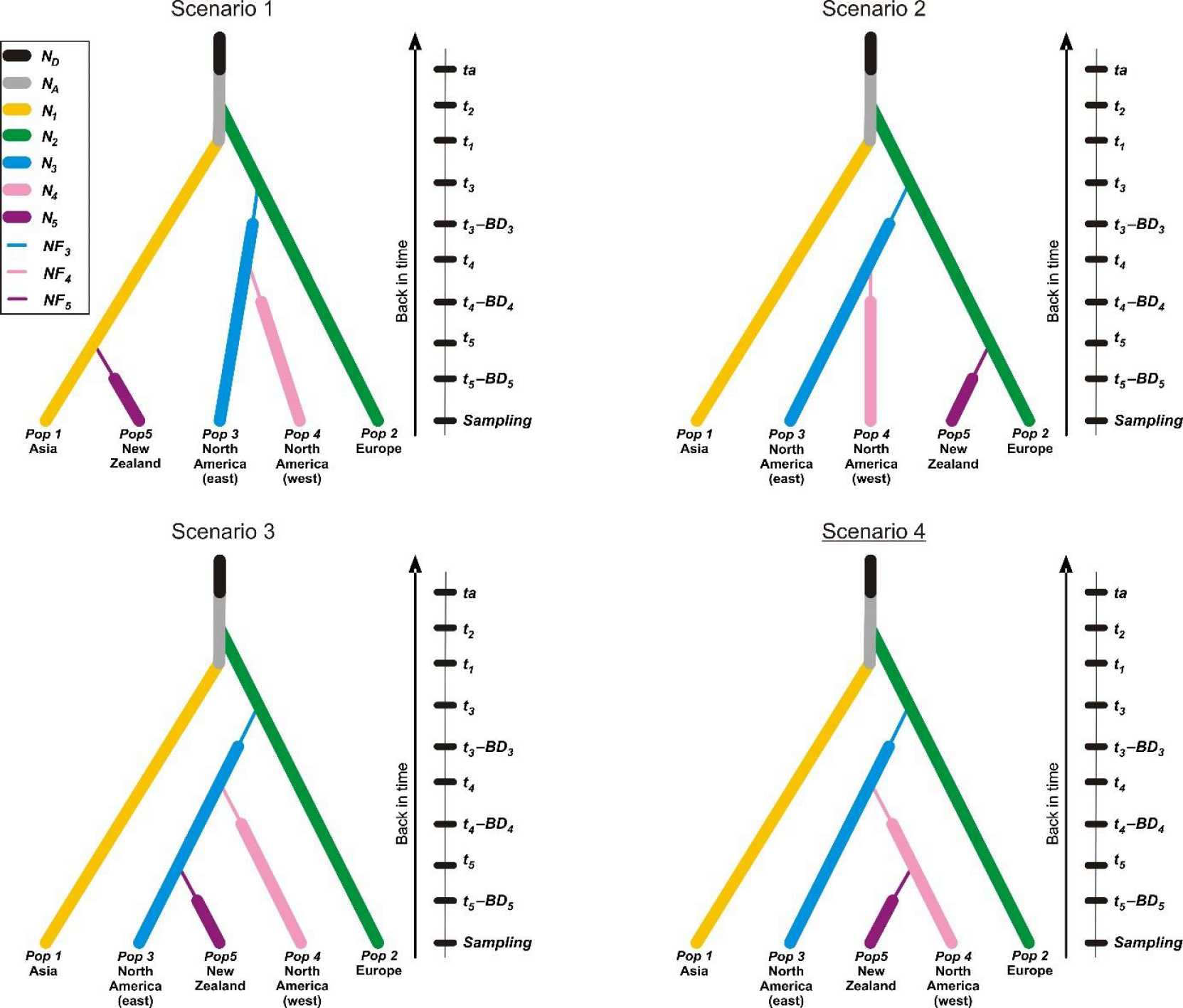
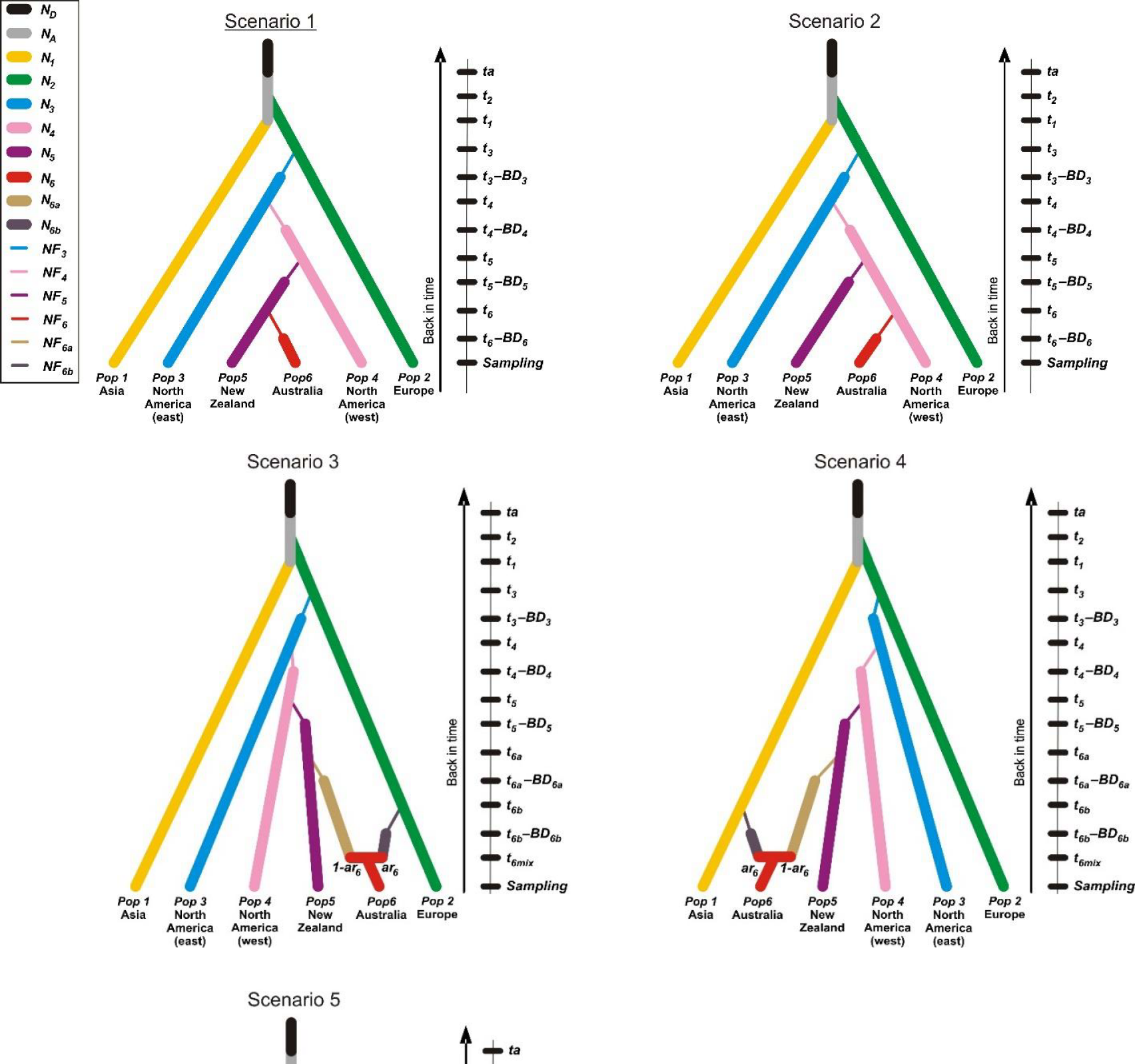
Schematic representation of each set of scenarios used in the ABC analyses to decipher the worldwide invasion routes of *Pieris rapae* (see also Table 1). Population numbers are as follows: 1 for Asia; 2 for Europe; 3 for North America (east); 4 for North America (west); 5 for New Zealand; 6 for Australia; 7 for North Africa; 8 for Russia (east). For each analysis, the name of the most likely scenario is underlined. Thin lines indicate bottlenecks. For parameters descriptions and priors see Table S5. Time is not to scale.

**Fig S4.**
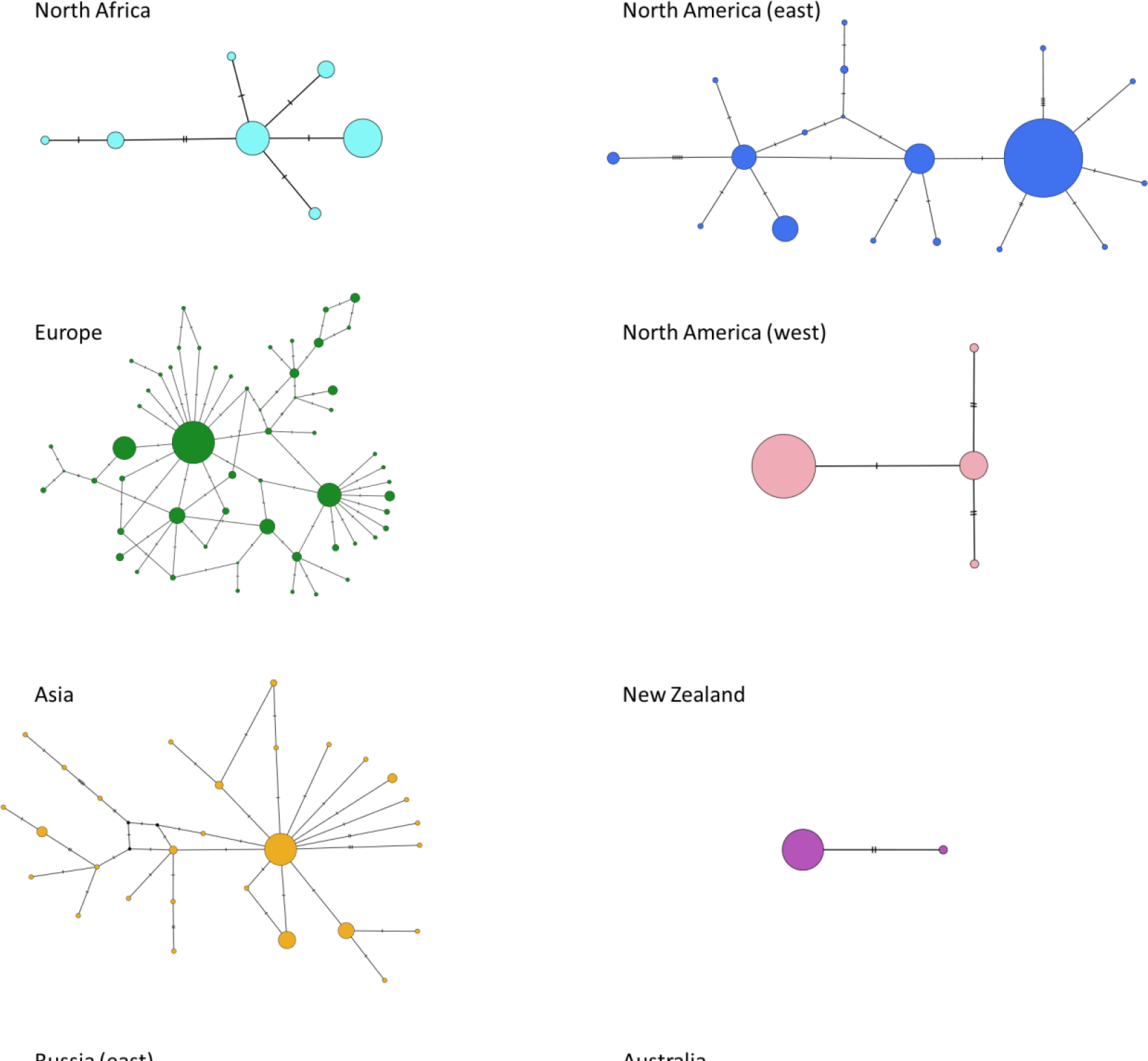
Median-joining haplotype networks for each population. Hash marks between haplotypes represent base changes (mutations).

**Fig S5.**
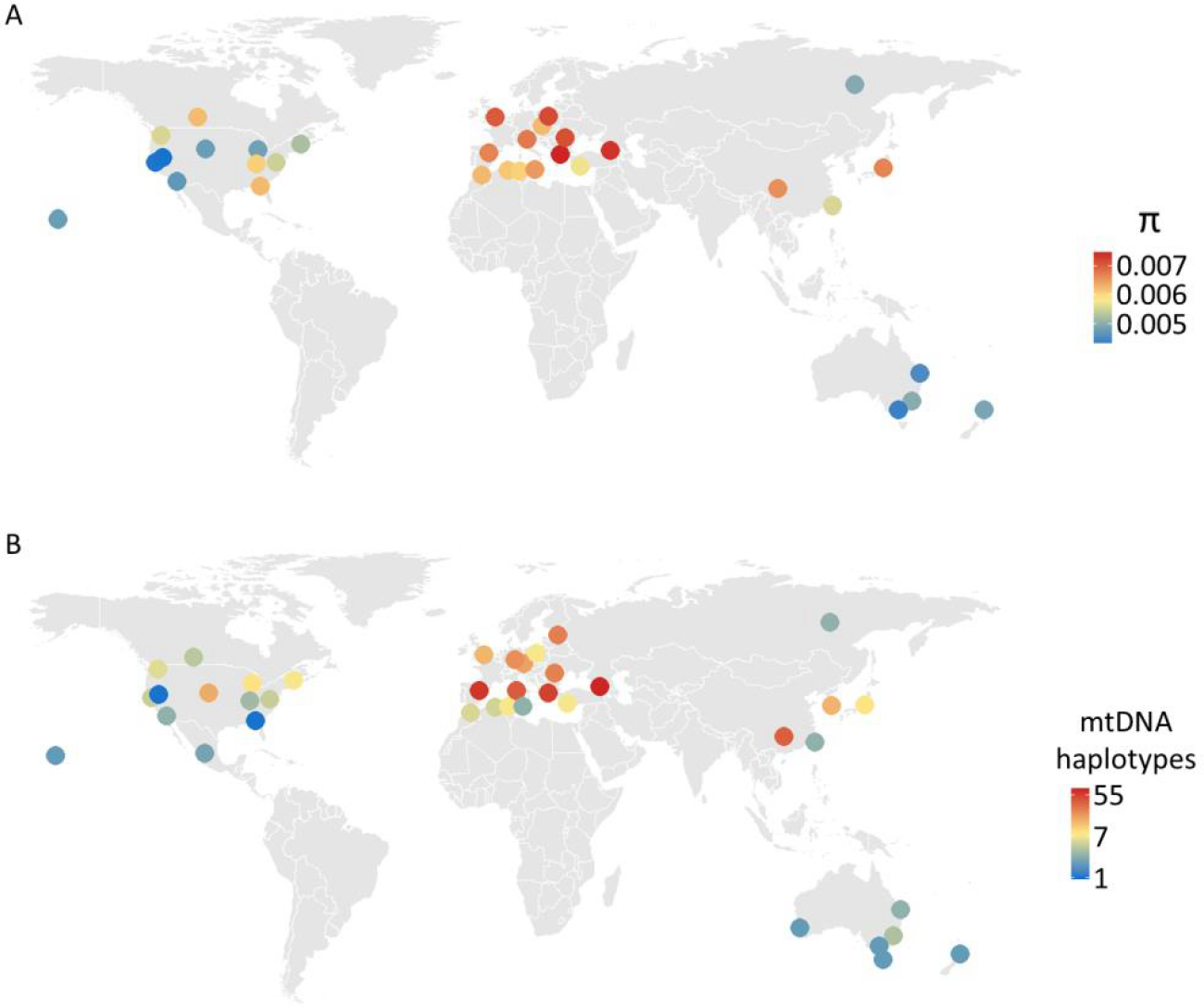
Global patterns of genetic diversity. **a,** Estimate of pairwise nucleotide diversity for each subpopulation based on autosomal ddRADseq data. **b**, mtDNA haplotype diversity estimated from rarefaction curves (note, colors are based on a log scale).

**Fig S6.**
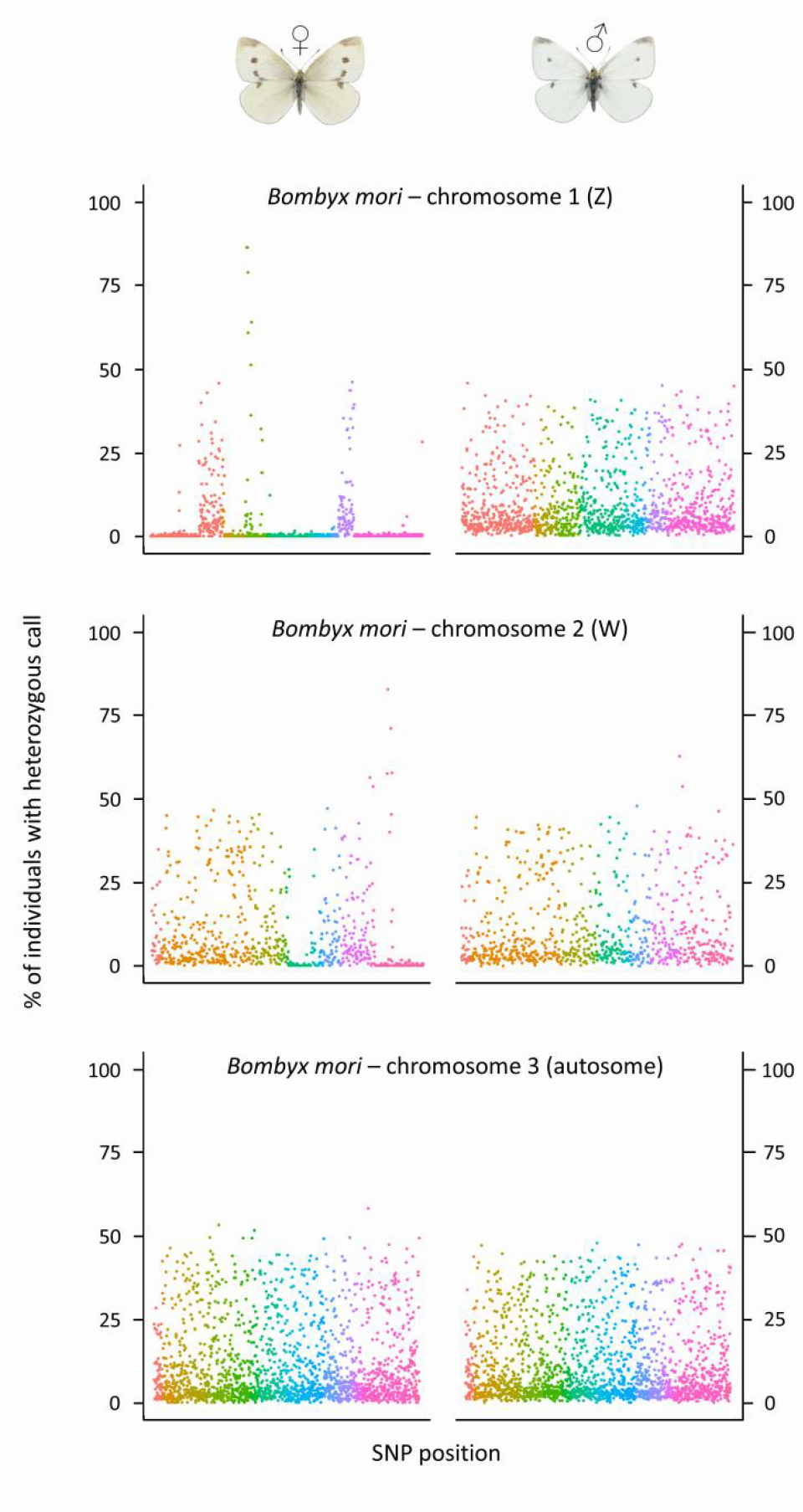
Percentage of individuals with heterozygous calls for each locus, plotted separately females and males. The location of each locus is based on its position within each *P. rapae* scaffold, with each *P. rapae* scaffold then ordered in each *B. mori* chromosome based on its homology to each *B. mori* scaffold (see Methods). Loci are colored by the *B. mori* scaffold to which they are associated. An autosome (chromosome 3) is plotted for reference and the pattern reflects those observed in other autosomes—no discernable difference in heterozygosity between males and females. Note, the W chromosome was not sequenced or assembled in the reference genome used in this study and is thus likely to be made up of portions of other chromosomes, including the Z (regions with no heterozygosity in females).

*Video included as a supplementary file*

**Video S1.** Development of railroad lines in the United States from 1830-1972. Railroad line data were obtained from Atack, 2016^28^ and plotted by their date of operation. Note the competition of railroad lines connecting eastern and western US in 1872, a few years prior (1879) to when a small population originating from North America (east) was believed to be introduced to that exact region—North America (west) (i.e., central California).

